# Targeted p63 isoform switch corrects dominant mutations in AEC syndrome without disrupting epidermal homeostasis

**DOI:** 10.1101/2025.06.24.661308

**Authors:** Daniela Di Girolamo, Gloria Urciuoli, Stefano Sol, Marco Ferniani, Dario Antonini, Emanuela Marchese, Maria Angela De Stefano, Marijke P A Baltissen, Marian A. Ros, Jill Dixon, Smail Hadj-Rabia, Caterina Missero

## Abstract

The transcription factor p63 is a master regulator of stratified epithelial development, and its disruption causes severe congenital defects affecting the skin, limbs, and craniofacial structures in both humans and mice. Among p63-related disorders, Ankyloblepharon-Ectodermal Defects-Cleft Lip/Palate (AEC) syndrome is caused by dominant mutations primarily affecting the Sterile Alpha Motif (SAM) domain and the Transactivation Inhibitory Domain (TID) of the *TP63* gene, which are unique to the p63α isoform. These mutations promote protein aggregation and transcriptional dysregulation, ultimately leading to debilitating skin erosions, suggesting that isoform-specific strategies could be therapeutically relevant.

To explore a therapeutic strategy based on isoform switching, we generated a conditional mouse model with deletion of exon 13, resulting in replacement of p63α by the shorter p63β isoform, which is expressed in the skin at lower levels. Although we found that p63α is required for limb and palate development, p63β proved sufficient to support epidermal formation, postnatal skin homeostasis, and wound healing. At the molecular level, the switch from p63α to p63β preserved chromatin binding and global transcriptional programs in keratinocytes.

We next used genome editing to delete exon 13 in human primary keratinocytes, inducing a switch from p63α to p63β. This isoform switch maintained normal proliferation and global gene expression. Importantly, p63β expression in AEC patient-derived keratinocytes rescued protein aggregation, restored mechanical integrity, and normalized epidermal gene expression. Together, these findings demonstrate that p63β can functionally compensate for p63α in the skin and establish and indicate that isoform switching could offer a new treatment option for AEC syndrome.

## Introduction

The transcription factor p63 (encoded by the *TP63* gene in humans, and the *Trp63* gene in mice) is a critical regulator of ectoderm-derived tissues, including the epidermis, craniofacial structures, and limbs (Mills, Zheng et al., 1999, Yang, Schweitzer et al., 1999). A member of the p53 family, *TP63* encodes several isoforms generated through alternative promoter usage (TA and ΔN) and C-terminal splicing (reviewed in (Di Girolamo et al., 2025)). While TAp63 plays a key role in the DNA damage response in oocytes by orchestrating apoptosis and preserving genomic integrity (Deutsch, Zielonka et al., 2011, Suh, Yang et al., 2006), ΔNp63 is the predominant isoform in stratified epithelia, where it is essential for epidermal commitment and stratification during embryonic development (reviewed in (Di Girolamo et al., 2025)). ΔNp63 expression marks the surface ectoderm early in embryogenesis and later becomes restricted to basal proliferative keratinocytes. Genetic ablation of *Trp63* or specifically of the ΔNp63 isoform disrupts epidermal development, leading to a disorganized, hypoplastic epidermis and absence of skin appendage (Mills et al., 1999, Romano, Smalley et al., 2012, Yang et al., 1999). These defects are also accompanied by craniofacial malformations and limb truncations, underscoring the essential role of p63 in ectodermal development. At the molecular level, ΔNp63 positively regulates a broad set of genes involved in keratinocyte proliferation and differentiation, as well as genes encoding components of desmosomes, hemidesmosomes, and the basement membrane, while simultaneously repressing genes associated with non-stratified epithelia (reviewed in (Di Girolamo et al., 2025)).

All p63 proteins share a conserved DNA-binding domain (DBD) and oligomerization domain (OD) but differ at the C-terminus due to alternative splicing. While the functions of the N-terminal p63 isoforms (TA and ΔN) are well characterized, the roles of the C-terminal variants are less clearly understood. Among these, the two predominant isoforms are p63α and p63β. The primary isoform in stratified epithelia is p63α that contains a 130-amino-acid C-terminal extension comprising the Sterile Alpha Motif (SAM) and the Transactivation Inhibitory Domain (TID), and accounts for approximately 80% of total p63 protein in the epidermis. In contrast, the p63β isoform lacks both the SAM and TID domains and instead contains a unique stretch of five alternative amino acids. It is the second most abundant isoform in the human adult epidermis, accounting for approximately 20% of total p63 expression (Marshall, Beeler et al., 2021, Rizzo, Romano et al., 2015). The SAM domain is a highly conserved protein–protein interaction module, present in the α isoforms of p63 and p73, but absent in p53. It is found in p63/p73 homologs across a wide range of species—from *C. elegans* and the American lobster to mammals—suggesting an evolutionarily preserved function (Ou, Lohr et al., 2007, Polinski, Zimin et al., 2021), although its precise role in epithelial tissues remains unclear. The TID is best characterized in oocytes, where it keeps TAp63α in an inactive conformation under physiological conditions and prevents premature activation of its pro-apoptotic function (Deutsch et al., 2011). In contrast, the role of the TID in ΔNp63α is less well defined, as this isoform is generally considered constitutively active— acting as both a transcriptional activator and repressor—without requiring the conformational activation observed in TAp63α. Notably, ΔNp63α-mediated repression involves interaction with histone deacetylases, and the TID is required for binding to HDAC1 and HDAC2, facilitating the formation of an active transcriptional repressor complex (LeBoeuf, Terrell et al., 2010, Ramsey, He et al., 2011).

Importantly, most pathogenic mutations causing AEC syndrome (OMIM #106260) affect the SAM and TID domains of the *TP63* gene (reviewed in (Di Girolamo et al., 2025)), highlighting the need to better understand the specific roles of p63α isoforms in development and disease. These dominant-negative mutations impair the regulatory functions of p63α, resulting in a severe developmental disorder characterized by ectodermal dysplasia, skin erosions, ankyloblepharon, cleft palate, and limb anomalies (Hay & Wells, 1976, McGrath, McMillan et al., 1997). Of all clinical features, recurrent scalp erosions are often the most debilitating and difficult to manage, frequently leading to secondary infections (Aberdam, Roux et al., 2020, Julapalli, Scher et al., 2009, McGrath, Duijf et al., 2001). At present, treatment relies solely on wound care, with no curative options available. We previously showed that AEC-associated mutations primarily affect the p63α isoform, disrupting its structural stability and leading to protein aggregation and impaired transcriptional activity (Russo, et al., 2018). These aggregates also exert dominant-negative effects by sequestering wild-type p63, causing widespread gene dysregulation. Notably, p63 variants that abolish aggregation of mutant proteins can restore transcriptional activity in reporter assays, as well as rescue epidermal gene expression in a human fibroblast-to-keratinocyte conversion model, implying that aggregation plays a central role in the molecular pathology of AEC syndrome.

p63β is co-expressed with the predominant ΔNp63α isoform in basal epithelial compartments, yet its specific role in epidermal development and homeostasis remains insufficiently defined. Previous studies have shown that complete loss of the C-terminal p63 isoforms—including both α and β—leads to developmental defects that, although milder, recapitulate key features of the full p63 knockout, such as epidermal hypoplasia, reduced progenitor cell proliferation, and limb malformations (Suzuki, Sahu et al., 2015). However, whether exclusive expression of p63β is sufficient to sustain epidermal function and compensate for the loss of p63α has not been directly tested.

Here, we investigated whether redirecting isoform expression toward p63β could preserve p63 function. Given that mutations in the SAM and TID domains destabilize p63α and promote protein misfolding and aggregation (Russo et al., 2018), we hypothesized that removing this region could restore p63 activity. To test this, we generated a conditional mouse model and CRISPR-edited human keratinocytes lacking exon 13, resulting in the loss of p63α and compensatory upregulation of p63β, without altering total p63 transcript or protein levels. This approach allowed us to assess whether p63β can functionally replace p63α in maintaining epidermal integrity and preventing disease-related phenotypes.

## Results

### Selective deletion of the p63α C-terminus reveals isoform-specific developmental roles

To investigate the specific function of the p63α C-terminus in development and disease, we generated conditional mutant mice (p63Δ13^fl/fl^) carrying a targeted deletion of *Trp63* exon 13, which is unique to the p63α isoform (Figure 1A and Suppl. Figure 1A). Exon 13 deletion was validated *in vitro* using primary keratinocytes derived from wild-type (p63), p63Δ13^fl/fl^ or p63Δ13^+/fl^ mice. Cells were infected with an adenovirus expressing Cre recombinase or a GFP control (Suppl. Figure 1B). Western blot analysis revealed a marked reduction in p63α levels, accompanied by a corresponding increase in p63β in Cre-transduced p63Δ13^fl/+^ cells. In Cre-transduced p63Δ13^fl/fl^ cells, p63α was completely absent and replaced by p63β (Suppl. Figure 1C).

**Figure 1.**
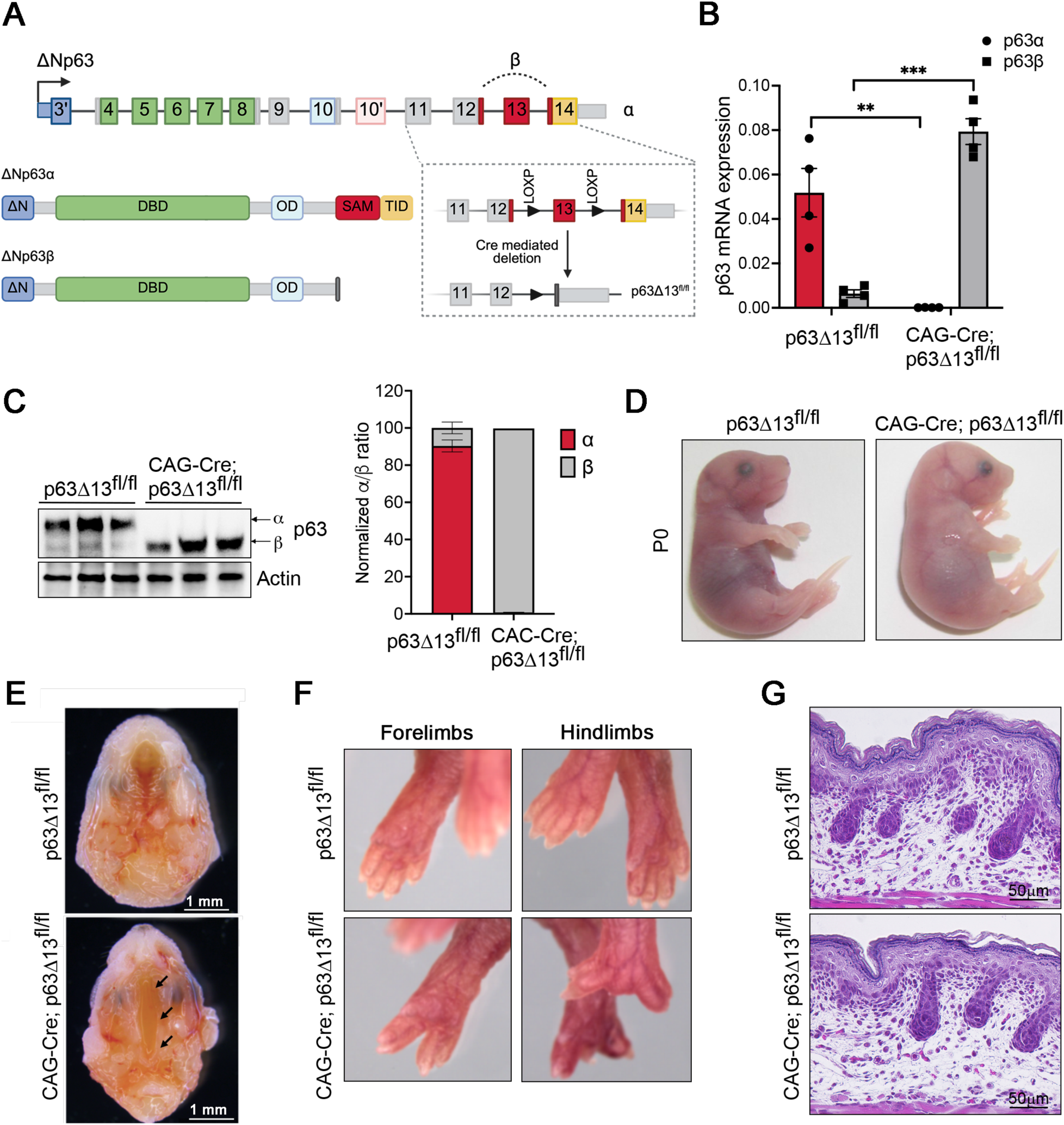
**Generation and characterization of CAG-Cre;p63Δ13^fl/fl^ mice.** A. *TP63* gene structure and strategy used to generate conditional p63Δ13^fl/fl^ mouse model. LOXP sites are flanking the wild type exon 13. Cre mediated deletion generates p63Δ13 mice. Exons of the ΔNp63 are indicated. Bottom left: ΔNp63α and ΔNp63β protein isoforms are shown with the indicated domains. In dark grey the short carboxyl terminal domain (5 amino acids) of the ΔNp63β isoform generated by alternative splicing of exon 13. Created with BioRender.com. B. Real Time RT-PCR performed on E18.5 epidermis RNA (n = 4 embryos/genotype). Data are the mean ± SD. Two-tailed paired Student’s t-test: **P < 0.005, ***P < 0.0005. C. Western blot assessing the levels of ΔNp63α and ΔNp63β proteins of E18.5 epidermis from p63Δ13^fl/fl^ and CAG-Cre; p63Δ13^fl/fl^ embryos (n = 3 embryos/genotype). Actin was used to normalize samples. Quantification of the ΔNp63α/ΔNp63β ratio normalised to Actin is shown on the right. D. Representative image of p63Δ13^fl/fl^ and CAG-Cre; p63Δ13^fl/fl^ mice at P0, showing illustrating normal skin appearance in the absence of p63α. E. Representative image of cleft left palate in CAG-Cre; p63Δ13^fl/fl^ observed at full penetrance compared to control p63Δ13^fl/fl^ mice at E18.5. The arrows point to the palatal cleft. Scale bar: 1mm. F. Representative image of limb malformations (Forelimbs and Hindlimbs) in CAG-Cre; p63Δ13^fl/fl^ compared to control p63Δ13^fl/fl^ mice at P0. G. Representative image of Hematoxylin and Eosin (H&E) staining on skin sections from p63Δ13^fl/fl^ and CAG-Cre; p63Δ13^fl/fl^ mice at P0. Scale bar: 50 μm.

p63Δ13^fl/fl^ mice were than crossed with CAG-Cre mice, which ubiquitously express Cre recombinase from early embryonic stages (Nagy, 2000), to induce deletion of exon 13 *in vivo*. This strategy led to loss of p63α and robust expression of the p63β isoform, which naturally lacks exon 13 (Figure 1A). Heterozygous mice were viable and fertile, and males were crossed with p63Δ13^fl/fl^ females to generate homozygous offspring. At E18.5, genotypes were distributed near Mendelian expectations: 18.3% CAG-Cre; p63Δ13^fl/fl^, 61% CAG-Cre; p63Δ13^+/fl^, and 20.7% p63Δ13 ^fl/fl^ controls (n = 82). However, CAG-Cre; p63Δ13^fl/fl^ mice died within hours from birth due to cleft palate. Using isoform-specific primers to distinguish p63α from p63β, Real Time (RT)-qPCR analysis confirmed a marked accumulation of p63β mRNA in the epidermis of CAG-Cre; p63Δ13^fl/fl^ at the expense of p63α (Figure 1B). Sequencing of a PCR-amplified cDNA fragment spanning exons 12 to 14 confirmed correct splicing between exon 12 and exon 14, with no evidence of aberrant splicing and proper switching to the p63β reading frame (Suppl. Figure 1D). Western blot analysis confirmed the complete loss of the ΔNp63α protein, accompanied by a corresponding increase in a lower molecular weight band consistent with ΔNp63β (Figure 1C). At birth, CAG-Cre; p63Δ13^fl/fl^ mutants consistently displayed a wide cleft of the secondary palate without involvement of the lip, primary palate or other cranial or mandibular structures (Figure 1D,E). In addition, they displayed fully penetrant limb malformations, including ectrodactyly and digit fusion (Figure 1D,F and see also below). Notably, the skin of mutant CAG-Cre; p63Δ13^fl/fl^ mice was indistinguishable from that of wild-type mice (Figure 1D,G). To our knowledge, this is the first p63 mouse model in which limb and palate malformations occur in the absence of overt skin abnormalities.

### Selective loss of p63α impairs oral epithelial organization and distal limb morphogenesis

To investigate the role of the p63α isoform in early palatogenesis, we analyzed secondary palate development in CAG-Cre; p63Δ13^fl/fl^ embryos. Unlike *Trp63*-null mice, which form only rudimentary palatal processes (Mills et al., 1999, Yang et al., 1999), CAG-Cre; p63Δ13^fl/fl^ embryos at E13.5 developed palatal shelves similar in size and morphology to those of p63Δ13^fl/fl^ controls, extending downwards on either side of the tongue (Figure 2A, upper panels). However, aberrant fusion of the oral epithelium was already evident at this stage, particularly in the region adjacent to the developing molars. By E14.5, palatal shelves in both control and mutant embryos had elevated above the tongue. However, in mutants they remained widely spaced and failed to contact at the midline (Figure 2A, middle panels). Additional aberrant epithelial fusions were observed between the palate and the lower oral epithelium, as well as between the tongue and the oral floor, likely interfering with proper shelf reorientation. By E15.5, wild-type embryos had completed palatal fusion and the midline epithelial seam had resolved, whereas in mutants the palatal shelves remained widely separated, failing to meet in the midline (Figure 2A, lower panels).

**Figure 2.**
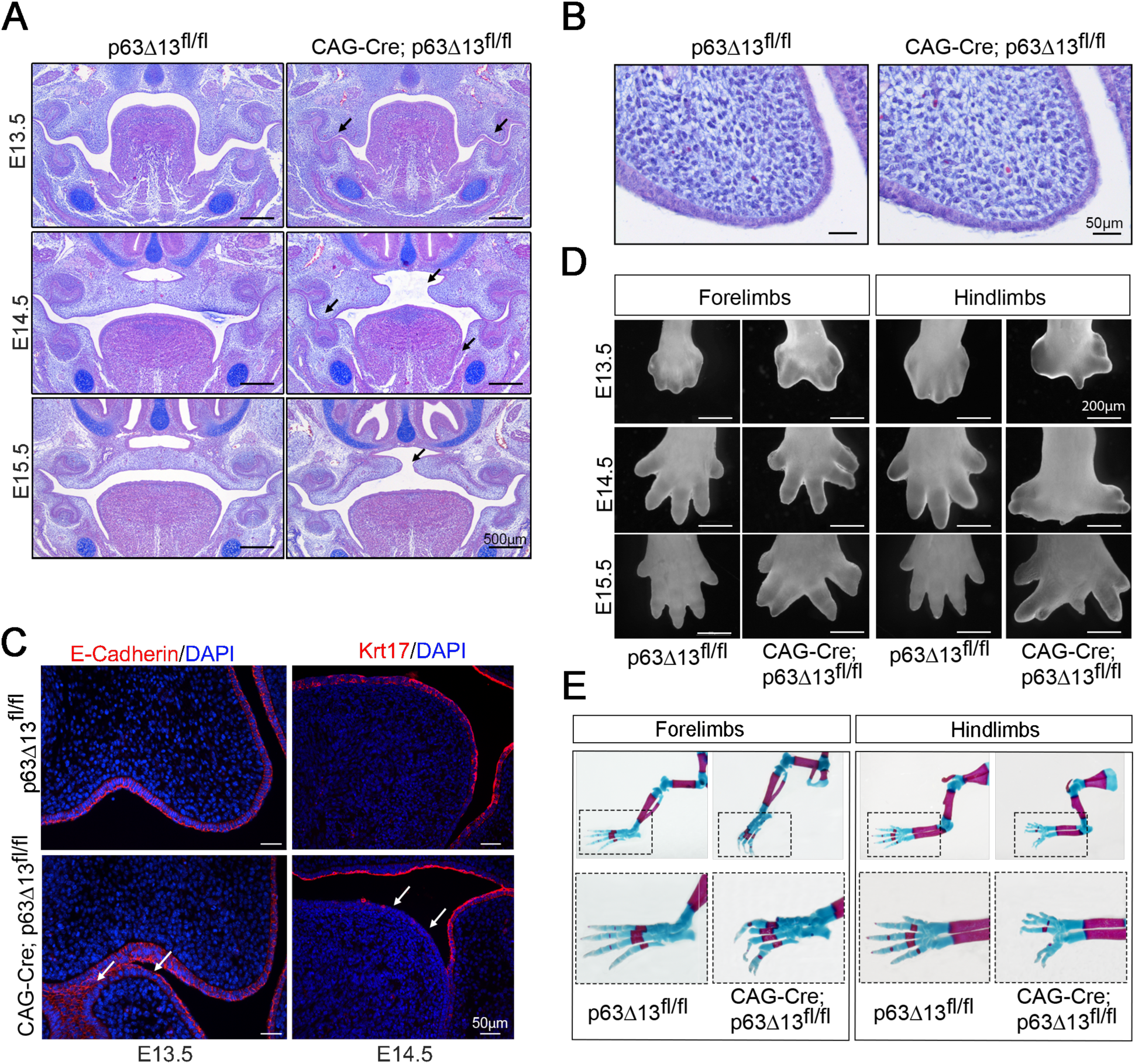
**Palate and limb characterization of CAG-Cre; p63Δ13^fl/fl^ mice.** A. Representative image of H&E staining of coronal palate sections of p63Δ13^fl/fl^ and CAG-Cre; p63Δ13^fl/fl^ mice at the indicated embryonic stages. Arrows point to epithelium fusion and separation of the palatal shelves in the mutant embryo. Scale bar: 500 μm. B. A closer view of E13.5 coronal anterior sections reveals that the epithelium of mutant palatal shelves is less organized compared to controls and the periderm is detaching from the surface ectoderm. Scale bar: 50 μm. C. Representative image of immunofluorescence analysis of palatal sections from p63Δ13^fl/fl^ *vs* CAG-Cre; p63Δ13^fl/fl^ embryos at E13.5 (left panels) and E14.5 (right panels). Sections were stained for E-Cadherin (red) and DAPI (blue, left panels), or for Keratin 17 (Krt17, red) and DAPI (blue, right panels). Arrows indicate epithelial defects in CAG-Cre; p63Δ13^fl/fl^ embryos, including aberrant epithelial fusions (E-Cadherin) and reduced peridermal Krt17 expression, compared to controls. Scale bar: 50 μm. D. Morphological analysis of limb development in p63Δ13^fl/fl^ and CAG-Cre; p63Δ13^fl/fl^ embryos at the indicated embryonic stages. CAG-Cre; p63Δ13^fl/fl^ embryos exhibit progressive malformations affecting the autopod, including central digit separation, ectrodactyly, and digit fusion, which are more pronounced in the hindlimbs. Scale bar: 200 μm. E. Representative image of skeletal analysis of forelimbs and hindlimbs in p63Δ13^fl/fl^ and CAG-Cre; p63Δ13^fl/fl^ embryos at E18.5. Alcian Blue/Alizarin Red staining reveals normal proximal limb structures and autopod malformations in mutants, including digit separation and fusion.

At higher magnification, the surface ectoderm of wild-type embryos consisted of basal columnar epithelial cells covered by a flattened, continuous layer of tightly connected periderm cells. In contrast, the periderm in CAG-Cre; p63Δ13^fl/fl^ mice appeared less flattened and more discontinuous (Figure 2B). At sites of aberrant epithelial fusion, continuous layers of E-Cadherin-positive cells were detected in mutants (Figure 2C), indicating inappropriate maintenance of epithelial adhesion. Alterations in the peridermal layer were also evident at E14.5. Krt17-positive peridermal cells were virtually absent in most epithelial regions of the tongue in CAG-Cre; p63Δ13^fl/fl^ embryos, consistent with the fusion events observed between the tongue, palatal shelves, and oral epithelium (Figure 2C). These findings indicate that loss of the p63α C-terminus impairs palatal shelf fusion, likely by disrupting periderm integrity and epithelial adhesion, without affecting initial shelf outgrowth or elevation.

Similarly, limb defects in CAG-Cre; p63Δ13^fl/fl^ mice were less severe than those observed in p63-deficient mice, which display poorly developed forelimbs and complete absence of hindlimbs (Mills et al., 1999, Yang et al., 1999). Mutant mice exhibited normal development of the stylopod (humerus/femur) and zeugopod (radius/tibia and ulna/fibula), but showed defects in the autopod. At E13.5, prior the onset of interdigit apoptosis in wild-type embryos, the distal limb of CAG-Cre; p63Δ13^fl/fl^ embryos consistently displayed split hand-like malformation, which was more pronounced in the hindlimbs (Figure 2D). By E15.5, CAG-Cre; p63Δ13^fl/fl^ embryos exhibited a range of limb defects, including ectrodactyly, syndactyly, and, more rarely, polydactyly affecting entire digits or individual phalangeal segments. The central region of the distal limb was most severely affected, with hindlimbs displaying a more pronounced phenotype than forelimbs. In particular, the hindlimbs frequently showed severe hypoplasia or complete absence of the central digit (Figure 2D). Alcian-Alizarin staining of E18.5 embryos confirmed normal development of the stylopod and zeugopod in CAG-Cre; p63Δ13^fl/fl^ mutant embryos. In contrast, the autopod showed skeletal defects, including central digit separation and partially missing or fused digits and bones (Figure 2E). This phenotype closely resembles Split Hand and Foot Malformation type 4 (SHFM4), which is associated with mutations in either the DNA-binding or the C-terminal domain of p63 (reviewed in (Osterburg et al., 2021)). Given the divergent phenotypes observed in skin and limb development, we investigated whether the relative abundance of p63α and p63β differs across these tissues. Analysis of publicly available RNA-seq data revealed that p63β is expressed at much lower levels in the developing epidermis compared to adult skin, and is nearly undetectable in the developing limb (Suppl. Figure 2), supporting a predominant role for p63α during early morphogenesis, particularly in limb development.

Taken together, these findings demonstrate that the p63α C-terminus is essential for proper morphogenesis of the secondary palate and distal limbs, highlighting its critical role in embryonic development.

### Selective p63α loss reveals compensatory function of p63β in skin development

Proper epidermal development depends on p63, as both p63-null and C-terminally truncated mouse models exhibit severe defects in stratification, differentiation, and progenitor cell proliferation (Mills et al., 1999, Senoo, Pinto et al., 2007, Suzuki et al., 2015, Yang et al., 1999). In contrast, replacement of p63α with p63β in CAG-Cre; p63Δ13^fl/fl^ newborn mice resulted in normal skin morphology and histology (Figure 1D,G). Immunostaining with a p63α-specific antibody confirmed the absence of this isoform in the epidermis of CAG-Cre; p63Δ13^fl/fl^ at birth (P0), while pan-p63 staining indicated persistent expression in basal keratinocytes, indicating compensation by p63β (Suppl. Figure 3A). Importantly, basal keratinocyte proliferation, assessed by Ki67 staining, was comparable between mutant and control epidermis (Figure 3A).

**Figure 3.**
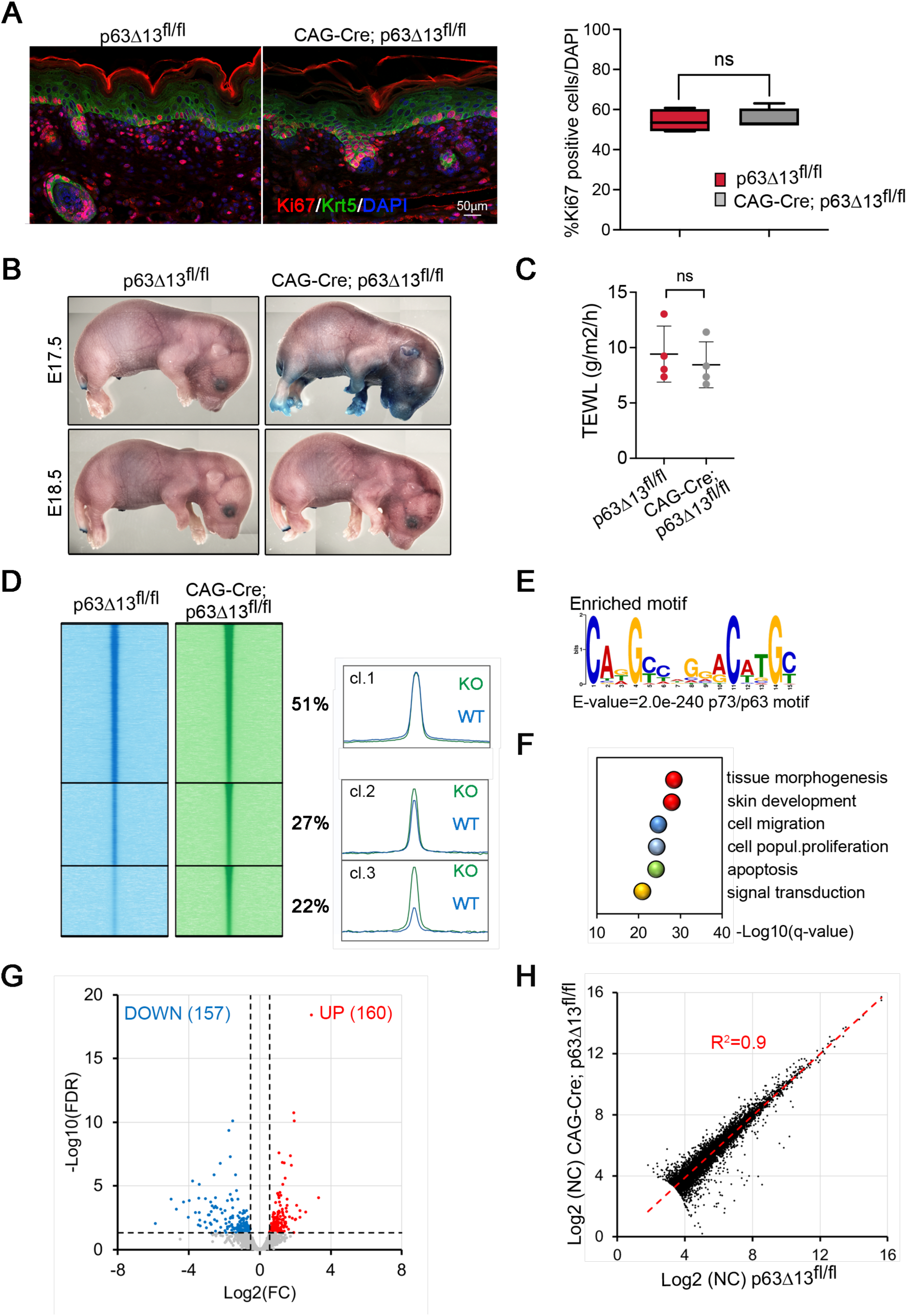
p**63β compensates for p63α loss with limited effects on epidermal chromatin binding and transcription.** A. Representative image of immunofluorescence analysis of Ki67 (red) and Krt5 (green) p63Δ13^fl/fl^ and CAG-Cre; p63Δ13^fl/fl^ epidermis at P0. Nuclei are stained with DAPI (blue). Right: quantification of Ki67^+^ cells relative to the total number of basal cells (n = 6 mice/genotype); ns = not statistically significant. Scale bar: 50 μm. B. Toluidine Blue exclusion assay at E17.5 and E18.5 shows transient delay in barrier formation in mutants. Images are stitched composites acquired using a stereomicroscope (Leica M205FA). C. TEWL measured at P0 shows no significant differences between genotypes (n = 4 mice/genotype). ns = not statistically significant. D. ChIP-seq analysis of p63 binding in P0 epidermis from control and mutant mice. High-confidence peaks (±5 kb from summit) were clustered by differential enrichment. 78% of peaks (clusters 1 and 2) are shared between genotypes, while 22% (cluster 3) are enriched in mutants. Heatmap was generated using ChAsE (Younesy et al., 2016). E. *De novo* motif analysis of the top 1,000 p63 binding regions common to both genotypes revealed enrichment of a canonical p63/p73 binding motif, with the indicated statistical significance. F. Gene Ontology (GO) biological processes enriched among genes associated with shared p63 binding regions in control and mutant epidermis. G. Volcano plot comparing gene expression in control p63Δ13^fl/fl^ and CAG-Cre; p63Δ13^fl/fl^ mutant E18.5 epidermis. Significantly upregulated (red) and downregulated (blue) genes are shown; gray: not significant. H. Linear regression analysis of RNA-seq data comparing gene expression between control p63Δ13^fl/fl^ and CAG-Cre; p63Δ13^fl/fl^ mutant epidermis at E18.5, demonstrating strong overall transcriptional similarity (R^2^=0.9).

Similarly, terminal differentiation appeared largely unaffected, as expression of canonical markers—keratin 5 (Krt5), keratin 14, (Krt14), keratin 10 (Krt10), involucrin (Ivl), loricrin (Lor), and filaggrin (Flg)— was comparable between mutants and controls (Figure 3A and Suppl. Figure 3A,B). However, reduced keratin 15 (Krt15) and induced keratin 6 (Krt6) expression indicated mild differentiation defects (Suppl. Figure 3C).

To assess whether the mild differentiation defects observed at birth reflected a more pronounced impairment during development, we assessed barrier acquisition during embryogenesis using Toluidine Blue dye exclusion. At E17.5, dye penetration into the ventral skin of mutant embryos indicated a mild delay in barrier formation (Figure 3B, upper panels), which was fully resolved by E18.5 (Figure 3B, and lower panels), in line with the nearly normal expression and localization of differentiation markers. Accordingly, Transepidermal Water Loss (TEWL) measurements at P0 showed no significant differences between genotypes (Figure 3C). These results indicate that switch from p63α to p63β causes only a transient delay in epidermal barrier maturation that is resolved by birth.

In the absence of overt histological abnormalities, we next asked whether switching from p63α to p63β alters the molecular landscape of the developing epidermis. To address this, we performed ChIP-seq analysis on P0 epidermis using a pan-p63 antibody. This approach allowed us to assess whether p63β, expressed in CAG-Cre; p63Δ13^fl/fl^ mice, binds the same regulatory elements as p63α *in vivo*, and to what extent isoform switching reshapes the p63 cistrome. Applying stringent peak-calling thresholds (see Methods), we found that the majority of p63-bound regions (76%) were shared between control (p63Δ13^fl/fl^) and mutant (CAG-Cre; p63Δ13^fl/fl^) samples (Figure 3D, clusters 1 and 2). These shared regions contained canonical p63 binding motifs (E-value = 2.0e-240) (Figure 3E) and were enriched for Gene Ontology (GO) biological processes related to tissue morphogenesis and skin development (Figure 3F). They were also associated with known p63 target genes (adjusted p-value = 5.053e-43; see Methods) (Figure 3F). Furthermore, 80% of these regions overlapped with ENCODE candidate cis-regulatory elements (cCREs) (Consortium, Moore et al., 2020) — a remarkably high proportion considering the limited annotation of regulatory elements in mouse tissues, and in particular, the paucity of data for the mouse epidermis.

Genomic regions preferentially bound in the mutant epidermis (Figure 3D, cluster 3) were similarly highly enriched for canonical p63 binding motifs (Suppl. Figure 3D), but showed markedly reduced overlap with ENCODE cCREs (41%). Interestingly, whereas regions bound in both genotypes frequently included multiple adjacent peaks within an 80 base pairs (bp) window, regions bound exclusively by p63β typically consisted of a single peak (Suppl. Figure 3E), This lack of peak redundancy suggests that p63β-specific sites may be less biologically meaningful. Given the redundancy of p63 binding across multiple regulatory elements for individual target genes, only a small number of genes were uniquely associated with p63β-specific binding regions. These genes showed no significant enrichment for known p63 targets or for targets of other transcription factors (data not shown). Collectively, these findings suggest that in the absence of p63α, p63β engages a subset of non-canonical regulatory elements with limited impact on core p63-driven transcriptional programs.

To assess the transcriptional consequences of p63α-to-p63β switching, we performed RNA-seq analysis on E18.5 embryonic epidermis (Figure 3G). Only 160 genes were significantly upregulated and 157 downregulated in mutants (Figure 3G)—a modest number relative to the widespread transcriptional changes reported in p63-null or AEC syndrome mouse models (Fan, Wang et al., 2018, Ferone, Thomason et al., 2012). In fact, linear regression revealed a high degree of similarity between control (p63Δ13^fl/fl^) and mutant (CAG-Cre; p63Δ13^fl/fl^) samples (R² = 0.9; Figure 3H). Among upregulated genes were *Trp63* itself, consistent with its established autoregulatory feedback loop (Antonini, Rossi et al., 2006, Antonini, Sirico et al., 2015), as well as canonical p63 targets such as *Itga3* and *Mdm2* (Suppl. Figure 3F). Conversely, downregulated genes were enriched for extracellular matrix and basement membrane components (e.g. *Col16a1*, *Col5a2*, *Eln*), and the stem cell-associated marker *Krt15*, suggesting a mild impairment in dermal–epidermal interactions and stem cell maintenance (Suppl. Figure 3F).

Integration of the RNA-seq and ChIP-seq datasets identified a limited subset of genes directly affected by the p63 isoform switching, with 145 genes, including Krt15 and Krt6, showing both differential expression and nearby p63 binding. This finding highlights the modest yet specific transcriptional consequences of replacing p63α with p63β. Overall, gene expression profiles were largely comparable between mutant and control epidermis, indicating that p63β can sustain the global transcriptional program of late epidermal development in the absence of p63α.

### p63β supports keratinocyte function and skin homeostasis in adult mice

To further characterize the effects of p63α loss and concomitant p63β expression on epidermal homeostasis at postnatal stages, we crossed p63Δ13^fl/fl^ mice with K14-Cre knock-in mice, in which Cre recombinase is inserted in the keratin 14 locus (Huelsken, Vogel et al., 2001). In this model, Cre is expressed from E17.5 onward in all stratified epithelia, enabling selective and complete deletion of exon 13 in the neonatal epidermis. Loss of p63α and concomitant induction of p63β in the neonatal epidermis were confirmed by Western blot and immunofluorescence using total p63 and p63α-specific antibodies (Figure 4A; Suppl. Figure 4A). As expected, K14-Cre; p63Δ13^fl/fl^ newborn mice were viable and displayed no limb or palate malformations, nor any overt skin defects (not shown). Adult K14-Cre; p63Δ13^fl/fl^ mice at two months of age were fertile, normal in size, and exhibited no visible skin defects (Figure 4B), nor did they develop any skin abnormalities at later stages. Consistently, histological analysis revealed no morphological abnormalities in the skin (Figure 4C).

**Figure 4.**
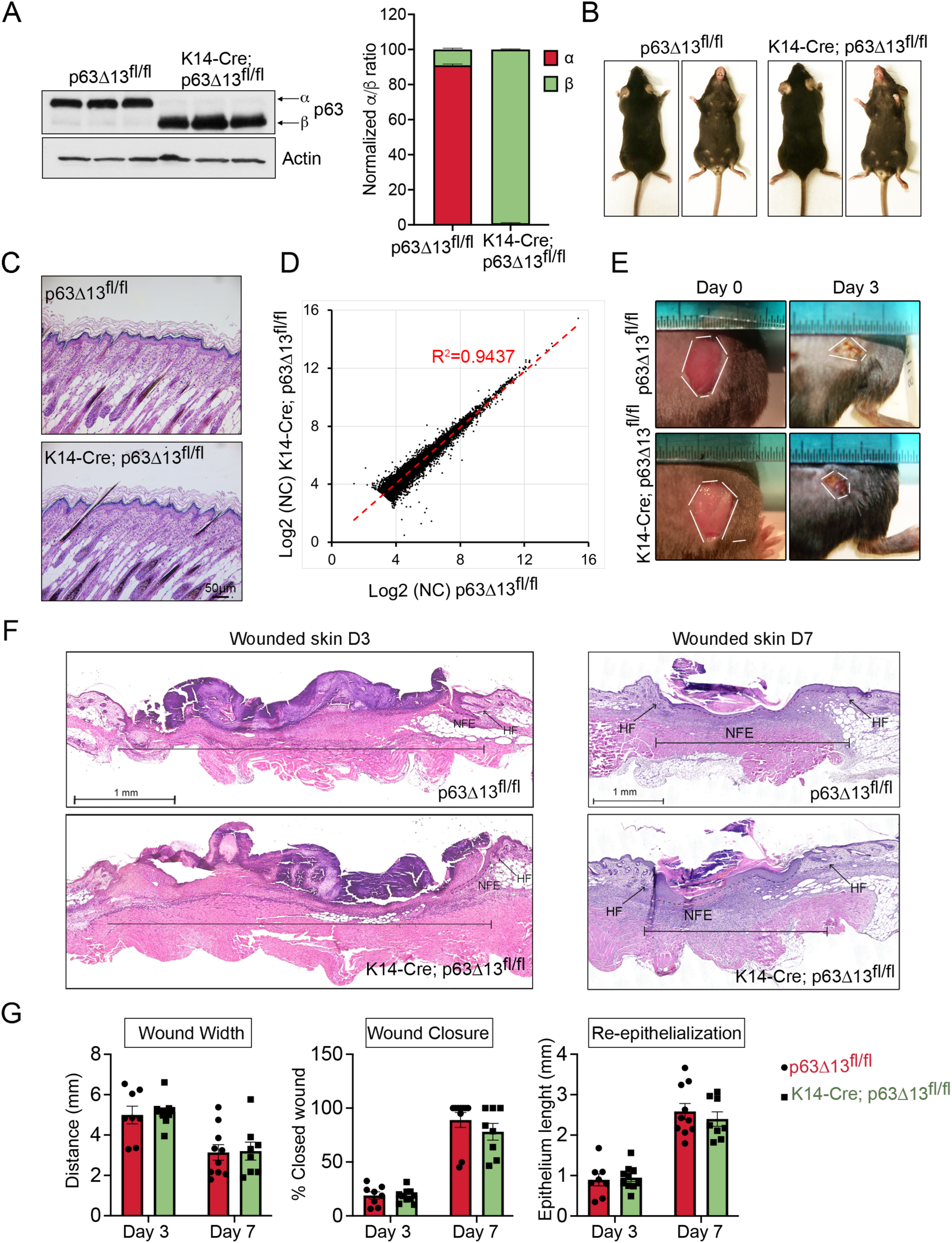
**Loss of p63α and induction of p63β under an epidermal specific promoter has no impact on epidermal homeostasis or wound healing.** A. Western blot analysis of ΔNp63α and ΔNp63β protein levels in P3 skin from control p63Δ13^fl/fl^ and K14-Cre; p63Δ13^fl/fl^ mutant mice (n = 3 mice/genotype). Actin was used to normalize samples. Right: quantification of the ΔNp63α/ΔNp63β ratio normalized to Actin. B. Representative image of adult (4 months old) p63Δ13^fl/fl^ and K14-Cre; p63Δ13^fl/fl^ mice showing comparable gross appearance. C. Representative image of H&E staining of P5 dorsal skin sections reveals normal epidermal architecture in both genotypes. Scale bar: 50 μm. D. Linear regression and Pearson correlation of RNA-seq data from primary mouse keratinocytes of p63Δ13^fl/fl^ and K14-Cre; p63Δ13^fl/fl^ mice shows strong transcriptional similarity (R^2^=0.9437). Normalized Count: NC. E. Representative macroscopic images of full-thickness excisional wounds at day 0 (D0) and day 3 (D3) post-injury. F. H&E-stained wound sections from p63Δ13^fl/fl^ and K14-Cre; p63Δ13^fl/fl^ mice at D3 and D7 (n = 8-10 mice per group per time point). HF: hair follicle; NFE: neo-formed epidermis. G. Morphometric analysis of wound width (left), percentage of wound closure (middle) and re-epithelization (right) at D3 and D7 (n = 8-10 mice per group per time point).

To assess whether the p63α-to-p63β isoform switch alters the transcriptome of keratinocytes, we performed RNA-seq analysis on P2 keratinocytes derived from K14-Cre; p63Δ13^fl/fl^ and p63Δ13^fl/fl^ control mice after 7 days in culture. *In vivo*, potential transcriptional alterations may be buffered by tissue-level homeostatic mechanisms and dermal-epidermal interactions. *In vitro* conditions subject keratinocytes to stress and remove the influence of their native microenvironment, which can amplify phenotypic differences. Despite this, transcriptomic analysis revealed no significant differences between genotypes, with a strong correlation (R² = 0.9437; Figure 4D), consistent with the absence of an overt skin phenotype (Figure 4B,C).

To further assess the functional role of p63β in maintaining skin integrity, we performed a wound healing assay in K14-Cre; p63Δ13^fl/fl^ and control mice. *In vivo* wound healing assays provide a physiologically relevant context to assess whether the loss of p63α impacts keratinocyte function during tissue regeneration and repair. Macroscopic measurements and morphometric analysis showed no significant differences in mean wound size between K14-Cre; p63Δ13^fl/fl^ and control mice at day 3 and day 7 post-wounding (Figure 4E,F). Quantification of the distance covered by the newly formed epidermis revealed no impairment in wound closure in mutant mice (Figure 4G). Similarly, wound epithelium length and wound width were comparable between genotypes, indicating that the switch from p63α to p63β does not affect re-epithelialization or wound contraction (Figure 4F,G).

In summary, even under stress of wound healing—which normally activates key pathways involved in cell migration, proliferation, and tissue repair—mutant keratinocytes displayed no functional impairments. These findings suggest that p63α and p63β play comparable roles in supporting keratinocyte function under both homeostatic and regenerative conditions.

### p63β compensates for p63α loss in human postnatal epidermal cells

We next examined the impact of expressing p63β in a human cellular context to explore whether the p63α-to-p63β switch could have therapeutic relevance for AEC syndrome. To assess whether human p63α and p63β isoforms similarly promote epidermal commitment, we firstly employed a transdifferentiation protocol that rapidly converts human dermal fibroblasts (HDFs) into keratinocyte-like cells (iKCs) via exogenous expression of ΔNp63α and Krüppel-like factor 4 (KLF4) (Chen, Mistry et al., 2014, Osterburg, Ferniani et al., 2023) (Suppl. Figure 4B). Human dermal fibroblasts isolated from neonatal skin (HDFn) or HDF-BJ-TERT (HDF-BJ) overexpressing p63β were successfully transdifferentiated into iKCs and expressed p63 target genes, including KRT14, KRT5, IRF6 and DSP, similarly to cells expressing p63α (Suppl. Figure 4C–F), indicating that p63β is also capable of inducing epidermal commitment in human cells.

To convert p63α into the p63β isoform in human primary keratinocytes (NHEK), we employed a dual small guide RNA (sgRNA) CRISPR/Cas9 strategy to delete exon 13 of the TP63 gene, thereby inducing isoform switching from p63α to p63β (Figure 5A). As shown in Figure 5B, bulk editing was highly efficient across four independent experiments. RNA-seq analysis confirmed these findings: in wild-type cells, 87% of *TP63* transcripts included exon 13, while 13% exhibited exon 13 skipping (Suppl. Figure 5A). In contrast, in the edited p63Δ13 keratinocyte pool, 94% of transcripts lacked exon 13. Western blot analysis further validated the isoform switch at the protein level in the bulk-edited keratinocytes (Figure 5C).

**Figure 5.**
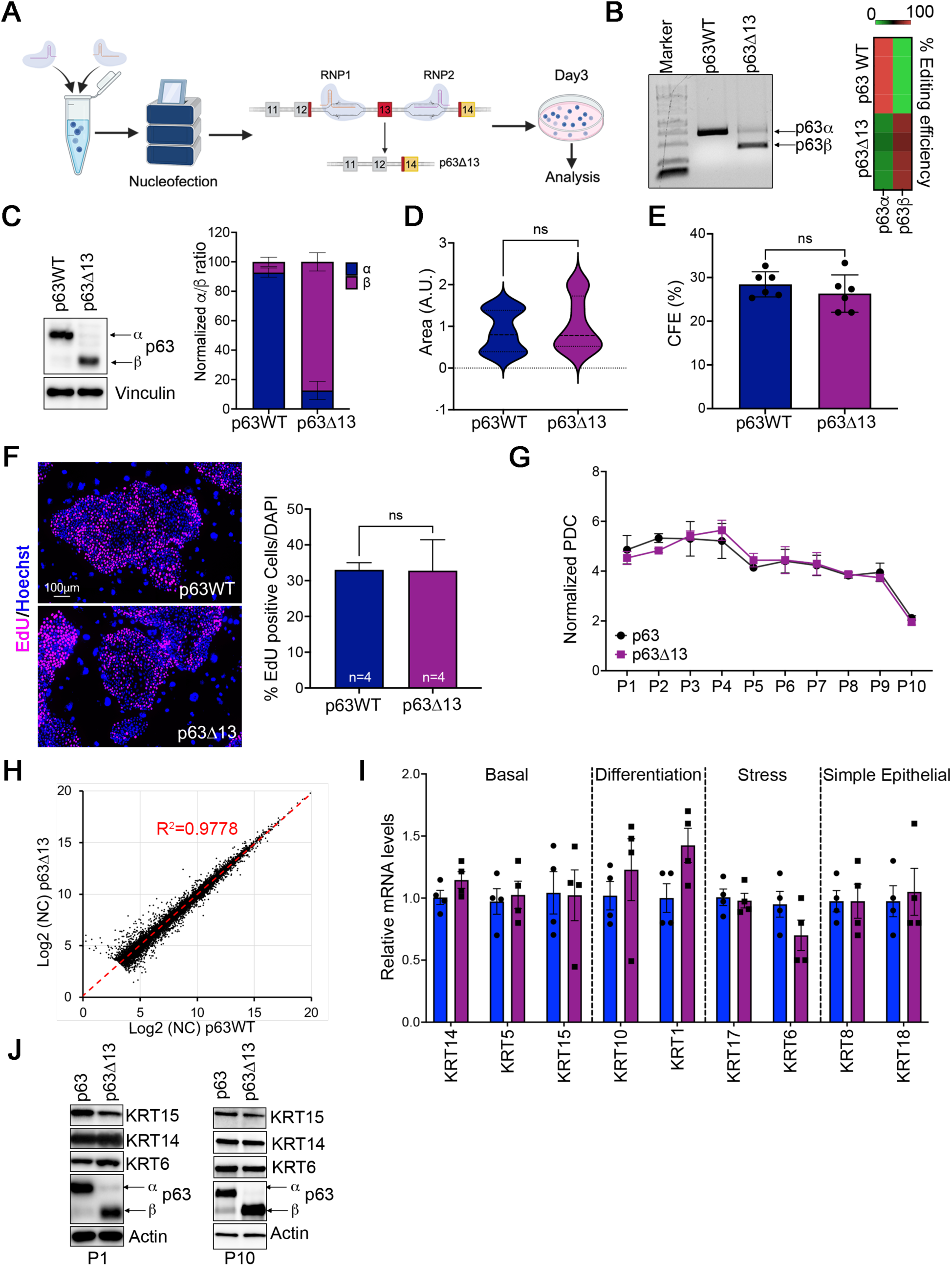
**CRISPR/Cas9-mediated deletion of exon 13 in human newborn keratinocytes preserves both short- and long-term proliferation and gene expression.** A. Scheme of the CRISPR/Cas9 editing strategy. RNP complexes targeting intronic regions flanking exon 13 (RNP1 and RNP2) were nucleofected into primary human keratinocytes. Exon 13 deletion via Non-Homologous End Joining (NHEJ) leads to p63α-to-p63β isoform switching. Cells were analyzed from day 3 post-nucleofection. Created with BioRender.com. B. PCR analysis of genomic DNA confirming exon 13 deletion in edited (p63Δ13) cells. Right: editing efficiency across four independent experiments (n=4). C. Western blot assessing the levels of ΔNp63α and ΔNp63β proteins from p63WT and p63Δ13 keratinocytes at day 3 post RNPs nucleofection. Vinculin was used to normalize samples. Bar graph shows the ΔNp63α/ΔNp63β ratio of quantified protein expression levels normalised to Vinculin (n = 4 independent experiments). D. Colony area measured in arbitrary units (AU) using ImageJ (n = 4 independent experiments). E. Colony-forming efficiency (CFE) expressed as percentage of colonies relative to plated cells (n = 6 independent experiments). F. EdU incorporation (purple) in p63WT and p63Δ13 keratinocytes. Nuclei stained with Hoechst (blue). Quantification of EdU⁺ cells over total nuclei (n = 4 independent experiments). Scale bar: 100 μm. G. Population doubling capacity (PDC) measured over passages using the formula: PD = 3.322 × log(N/N₀). No significant differences were observed (n = 4 independent experiments). H. Linear regression and Pearson correlation of RNA-seq data from p63WT and p63Δ13 keratinocytes shows strong transcriptional similarity (R^2^=0.9778). Normalized Count: NC. I. RT-qPCR for keratin genes in p63Δ13 vs p63WT keratinocytes (n = 4 independent experiments). J. Western blot analysis of KRT15, KRT14, KRT6, and p63 at different passages (P1 and P10) in p63WT and p63Δ13 cells. Actin was used to normalize samples.

To evaluate whether isoform switching affects the clonogenic potential of human keratinocytes, we assessed colony morphology and growth capacity in p63Δ13 and control NHEKs. Colony morphology appeared broadly similar between p63Δ13 and control NHEKs (Suppl. Figure 5B). Consistently, wild-type and edited NHEKs displayed comparable colony size and colony-forming efficiency (CFE) (Figure 5D,E).

To assess whether depletion of the p63α isoform affects the proliferative capacity of primary keratinocytes, we performed a 5-ethynyl-2’-deoxyuridine (EdU) incorporation assay. As shown in Figure 5F, p63Δ13 and control NHEKs exhibited comparable proliferation rates in bulk cultures. Consistently, no significant differences were observed in population doubling capacity (PDC) between the two groups, indicating that deletion of exon 13 does not alter the long-term proliferative capacity or replicative lifespan of human keratinocytes (Figure 5G). Moreover, the percentage of edited cells and the p63α/p63β protein ratio remained stable across cell passages (Suppl. Figure 5C–E), indicating that p63Δ13 NHEKs do not exhibit a proliferative advantage or disadvantage over time.

To investigate the impact of exon 13 deletion on gene expression profile of human primary keratinocytes, we performed RNA-seq analysis comparing p63Δ13 and control NHEKs. Transcriptomic analysis confirmed a successful isoform switch from p63α to p63β in edited samples, with p63β expression increasing from 13% in wild-type cells to 91% post-editing. Only minimal expression of unintended isoforms was detected—specifically, 3% of exon 11–14 splicing corresponding to the previously described p63δ isoform (Mangiulli, Valletti et al., 2009)— supporting the overall specificity and potential safety of the CRISPR/Cas9-mediated exon 13 deletion strategy (Suppl. Figure 5A). Interestingly, deletion of exon 13 in p63Δ13 NHEKs affected only a negligible number of genes, resulting in a highly significant positive correlation in global gene expression compared to control cells (R^2^ = 0.9778, Figure 5H). Accordingly, protein levels of several p63 target genes—including epidermal and simple epithelial keratins—remained unaffected by exon 13 deletion, even after multiple cell passages (Figure 5I-J). These findings demonstrate that p63β is sufficient to sustain the transcriptional and protein expression programs required for keratinocyte identity and homeostasis in the absence of p63α, further supporting the functional redundancy of these isoforms in epidermal cells.

### Replacement of mutant p63α with p63β restores functionality in human AEC keratinocytes

Given that autosomal dominant mutations in exons 13 and 14 of the *TP63* gene underlie AEC syndrome, we hypothesized that deleting exon 13 in the mutant allele would shift expression from p63α to p63β, thereby removing the domain most frequently affected by pathogenic mutations. To test this, we applied CRISPR/Cas9-mediated gene editing to two independent primary keratinocyte cultures derived from AEC patients harboring the p63-T533P and p63-I537T mutations (Figure 6A). Exon 13 deletion was achieved with high efficiency and minimal toxicity, comparable to editing outcomes in wild-type keratinocytes (Suppl. Figure 6A,B).

**Figure 6.**
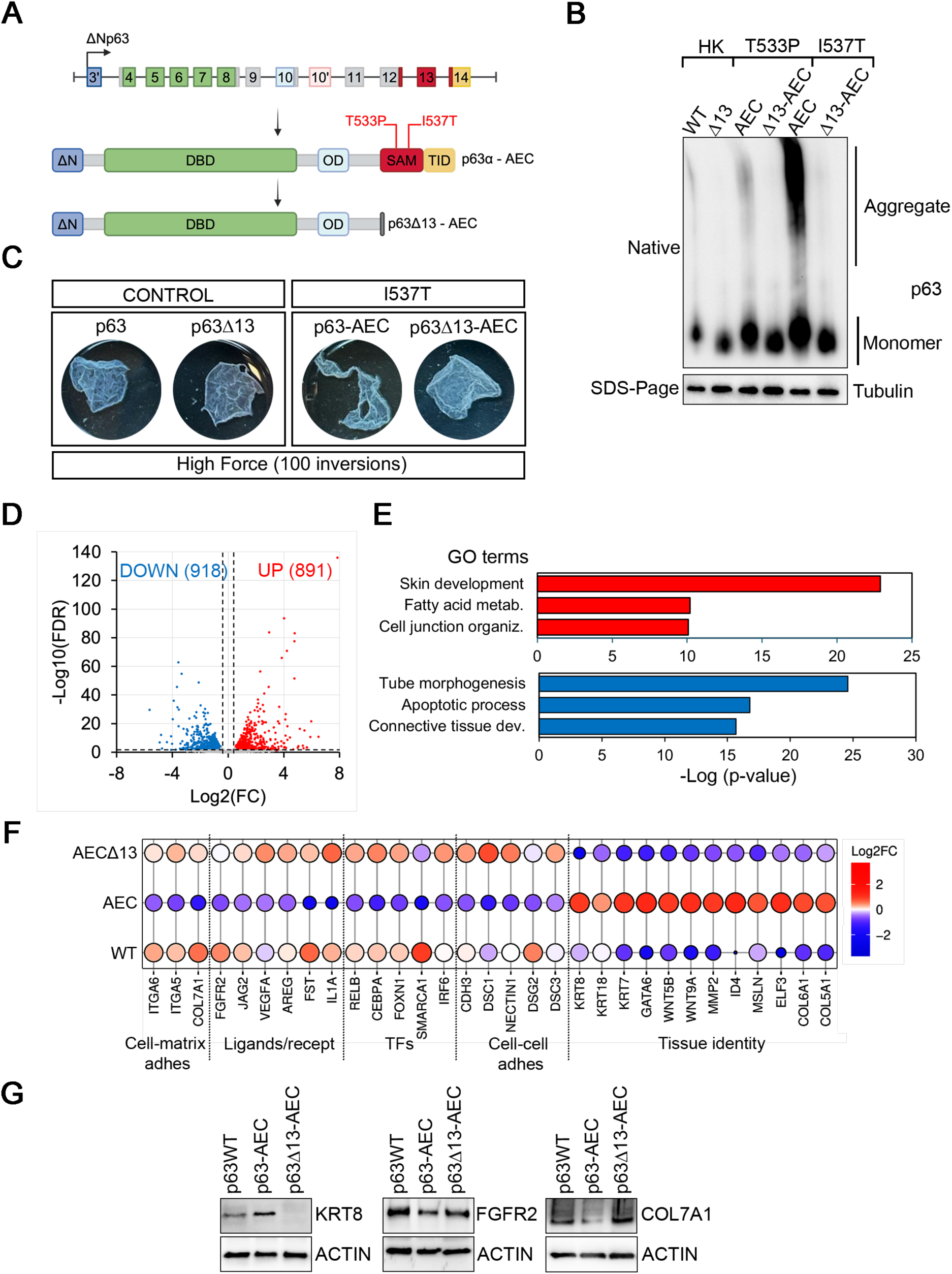
**Restoration of p63 function in AEC-derived keratinocytes by exon 13 deletion.** A. Scheme of AEC-associated mutations (T533P, I537T) and generation of p63Δ13-AEC keratinocytes by targeted deletion of exon 13. Created with BioRender.com. B. BN-PAGE (top) and SDS/PAGE (bottom) followed by Western blot for p63 and tubulin. Exon 13 deletion reduces aggregation in AEC-mutant keratinocytes and not in human adult keratinocytes (HK) (n = 3 independent experiments). C. Dispase-based epidermal sheet dissociation assay after mechanical stress (100 inversions). p63Δ13-AEC keratinocytes exhibit improved sheet integrity compared to p63-AEC (I537T) mutants, while no differences were observed between control NHEK and p63Δ13 keratinocytes. (n = 3 independent experiments). D. Volcano plot of differentially expressed genes in p63Δ13-AEC vs p63-AEC keratinocytes (n = 3 independent experiments). Significantly upregulated (red) and downregulated (blue) genes are shown; gray: not significant. E. GO enrichment analysis of biological processes associated with differentially expressed genes between p63Δ13-AEC and p63-AEC keratinocytes. Upregulated biological processes are in red and downregulated in blue. F. Ballon plot visualisation of selected p63 target genes across functional categories. Color intensity represents Log₂ fold change. G. Western blot analysis of selected p63 target genes in WT (NHEK), AEC, and Δ13-AEC keratinocytes. Actin was used to normalize samples.

We previously showed that AEC-associated *TP63* mutations induce protein misfolding and aggregation, leading to impaired p63 function (Russo et al., 2018). In line with these findings, patient-derived mutant keratinocytes exhibited prominent p63 aggregation (Figure 6B), suggesting that aggregation is a major contributor to the pathogenic loss of function. Strikingly, deletion of exon 13—and the resulting isoform switch from p63α to p63β—abolished aggregation and restored protein solubility (Figure 6B). These findings underscore the critical contribution of the C-terminal α domain to protein aggregation propensity and suggest that its removal may restore protein homeostasis.

To further investigate the functional consequences of this rescue, we assessed resistance to mechanical stress in mutant keratinocytes, as p63 directly transactivates several desmosome genes and defective cell-cell adhesion has been reported in an AEC mouse model (Ferone et al., 2013). Epidermal sheets were generated from control NHEK (p63WT), p63β-expressing NHEK (p63Δ13), mutant AEC keratinocytes (p63AEC-I537T), and p63β-expressing mutant keratinocytes (p63Δ13-AEC). Sheets derived from p63AEC-I537T keratinocytes were markedly more fragile and prone to disintegration under mechanical stress compared to wild-type controls (Figure 6C). Remarkably, expression of p63β in the mutant background (p63Δ13-AEC) restored mechanical resilience to levels comparable with both p63WT and p63Δ13 cells. These findings demonstrate that exon 13 deletion and consequent isoform switching from p63α to p63β not only rescue protein solubility but also restore epithelial cohesion and mechanical integrity in AEC mutant keratinocytes (Figure 6C), highlighting a direct link between protein aggregation, desmosome function, and tissue fragility.

To determine whether exon 13 deletion also rescues transcriptional defects associated with AEC mutations, we performed RNA-seq analysis comparing mutant keratinocytes (p63-AEC-I537T) to their exon 13-deleted counterparts (p63Δ13-AEC-I537T). Exon 13 deletion had a broad impact on gene expression, with more than 1,800 genes differentially expressed relative to the mutant control (Figure 6D). This contrasts with the minimal transcriptomic changes observed when exon 13 was deleted in wild-type keratinocytes (Figure 5H), underscoring the mutation-specific corrective effect of isoform switching. Genes upregulated following exon 13 deletion in AEC keratinocytes were highly enriched for genes involved in skin development (Figure 6E),

consistent with restored p63 function. Among the upregulated genes were numerous canonical p63 targets involved in epidermal differentiation, adhesion, and structural integrity (Figure 6F). Conversely, several genes previously reported as aberrantly upregulated in AEC syndrome and/or in p63 deleted or depleted keratinocytes (De Rosa, Antonini et al., 2009, Ferone, Mollo et al., 2013, Ferone et al., 2012, Romano et al., 2012, Russo et al., 2018, Truong, Kretz et al., 2006, Zarnegar, Webster et al., 2012) were significantly downregulated upon exon 13 deletion (Figure 6F). Notably, most of these transcriptional changes occurred in the opposite direction relative to those observed in untreated AEC mutant keratinocytes versus wild type ones, indicating that conversion of p63α to p63β can effectively rescue the mutant transcriptional program (Figure 6F). Consistent with the transcriptomic findings, western blot analysis confirmed that protein levels of selected p63 target genes were restored following exon 13 deletion and the resulting switch from p63α to p63β (Figure 6G). Together, these findings demonstrate that deletion of exon 13 and isoform conversion from p63α to p63β not only rescues protein solubility and mechanical resilience in AEC mutant keratinocytes but also restores the expression of critical epidermal genes at both transcript and protein levels.

## Discussion

Despite the evolutionary conservation of the SAM and TID domains of ΔNp63α and their causal role in AEC syndrome, their specific functions in stratified epithelia remain poorly understood. Here, using genetically engineered mouse models and CRISPR-edited human keratinocytes, we demonstrate that deletion of exon 13 removes the aggregation-prone C-terminal region of p63α, effectively converting it into the p63β isoform. This isoform switch does not alter overall p63 protein levels but has distinct functional outcomes: while p63β retains the ability to sustain epidermal transcriptional programs and homeostasis, it fails to support proper palatal fusion and digit morphogenesis. This represents the first *in vivo* model in which p63-dependent defects in skin, palate, and limb development are uncoupled. In previously described mouse models, ablation of p63 or ΔNp63, or expression of mutant p63 variants, causes severe malformations in all three tissues (Ferone et al., 2013, Ferone et al., 2012, Lo Iacono, Mantero et al., 2008, Mills et al., 1999, Romano et al., 2012, Russo et al., 2018, Suzuki et al., 2015, Yang et al., 1999). Similarly, deletion of exon 12—which leads to premature truncation of both p63α and p63β— disrupts skin, limb, and palate development (Suzuki et al., 2015). In contrast, our p63Δ13 model, which selectively eliminates p63α while preserving p63β, maintains normal epidermal function and transcriptome but exhibits cleft palate and distal limb malformations. These findings demonstrate that p63β is sufficient to sustain epidermal development but cannot fully compensate for p63α function in the palate and limbs.

The molecular mechanisms underlying the tissue-specific requirement for p63α remain to be elucidated. Analysis of the developing limb indicates that while p63β is sufficient for formation of the stylopod and zeugopod, it fails to support proper autopod development. p63 is expressed in the apical ectodermal ridge (AER), a specialized stratified ectodermal structure that promotes limb outgrowth by sustaining mesenchymal proliferation through FGF8 signaling (Lo Iacono et al., 2008, Mills et al., 1999, Yang et al., 1999). It is plausible that p63α, but not p63β, is required for the transcriptional activation of key regulators of digit identity, such as Dlx5 and Dlx6 (Kouwenhoven, van Heeringen et al., 2010, Lo Iacono et al., 2008), and for cooperative interactions with other limb-specific transcription factors. The restriction of the phenotype to the autopod and its late onset suggest that p63α is dispensable for early AER specification but plays a crucial role in later stages of distal limb morphogenesis. The strong resemblance to defects observed in Dlx5/Dlx6 loss-of-function models (Kouwenhoven et al., 2010, Lo Iacono et al., 2008, Robledo, Rajan et al., 2002) supports this hypothesis, although other factors are likely involved given the broad regulatory roles of p63.

Cleft palate in p63Δ13 embryos is associated with aberrant intraoral adhesions likely due to defective periderm formation and resemble phenotypes caused by reduced p63 and Irf6 mutations (Richardson, Dixon et al., 2009, Richardson, Hammond et al., 2014, Thomason, Zhou et al., 2010). Previous studies have shown that periderm cells act to form a protective barrier preventing pathological adhesion between intimately apposed, adhesion-competent epithelia. Mosaic ablation of periderm using a genetic strategy resulted in intra-oral adhesions between the maxillary and mandibular processes. Although these adhesions were generally limited to the lateral regions of the oral cavity, in severe cases they prevented normal development of the palatal shelves resulting in cleft palate (Richardson et al., 2014). In humans, disruption of periderm formation and/or function underlies a series of birth defects that exhibit multiple inter-epithelial adhesions including popliteal pterygium, cocoon and Bartsocas Papas syndromes. Genetic analyses of these conditions have shown that IRF6, IKKA, SFN, RIPK4 and GRHL3, all of which are under the transcriptional control of p63, play a key role in periderm formation (Hammond, Dixon et al., 2019, Richardson, Mitchell et al., 2017, Richardson et al., 2014). Clarifying whether the SAM domain, the TID domain, or both are essential for proper digit and palate development will be critical to understanding the mechanistic basis of p63 function in these tissues. Despite over 25 years of research on p63, the precise role of the SAM domain remains elusive. By contrast, the TID domain has been more extensively studied, particularly in the context of TAp63α function in oocyte quality control (Deutsch et al., 2011). In this setting, the TID domain maintains TAp63α in a closed, inactive, dimeric conformation, preventing premature activation of its pro-apoptotic transcriptional program, which is only triggered in response to DNA damage (Deutsch et al., 2011, Osterburg et al., 2021). In line with this regulatory role, ubiquitous heterozygous deletion of exon 13, which encodes part of the TID domain, leads to infertility in female mice, presumably due to inappropriate activation of TAp63α in oocytes (Lena, Rossi et al., 2021). Determining the relative contribution of these domains to the activity of ΔNp63α in non-gonadal epithelia will provide crucial insights into isoform-specific regulation and the pathogenesis of developmental disorders.

While p63β expression is low in the surface ectoderm during mid-gestation and even lower in the limb ectoderm, it reaches approximately 20% of total p63 transcripts by birth and in cultured keratinocytes (this work and (Marshall et al., 2021, Sethi, Romano et al., 2015)). The function of p63β and the regulation of exon 13 splicing have remained elusive. Surprisingly, skin expressing only p63β appears entirely normal and exhibits no detectable phenotype, including during stress conditions such as wound healing. Together, these findings suggest that ΔNp63β retains the core transcriptional functions necessary for keratinocyte identity and stratification, despite lacking the C-terminal domains. In addition, these data indicate that both the SAM and the TID domains are dispensable in postnatal skin, and that the few amino acids uniquely present in the C-terminal domain of p63β do not significantly alter p63 function in the epidermis and its appendages. Mechanistically, ChIP-seq and RNA-seq analyses confirm that p63β maintains occupancy at canonical p63 target loci and drives expression of major epidermal genes. Interestingly, p63β binds more genomic sites than p63α, yet these additional peaks do not correlate with gene expression changes or enrich for specific GO categories. This suggests that the α C-terminal region may fine-tune chromatin interactions rather than being essential for basic enhancer binding.

From a therapeutic perspective, these findings are highly relevant. Mutations in exons 13– 14 of *TP63*, which underlie the majority of AEC syndrome cases, destabilize the C-terminal domain, promote protein aggregation, and ultimately compromise transcriptional activity (Russo et al., 2018). Leveraging evidence from the mouse model, we developed a CRISPR-Cas9 strategy to delete exon 13 in patient-derived keratinocytes, effectively converting mutant p63α into functional p63β. This targeted editing reduced protein aggregation, improved mechanical resilience, and restored the expression of multiple genes dysregulated in AEC keratinocytes. Using two sgRNAs flanking exon 13, we achieved >80% editing efficiency, selectively rescuing p63 function in AEC cells without impacting wild-type keratinocytes. This proof-of-concept highlights isoform switching as a potential therapeutic strategy to restore skin integrity in AEC, with broad applicability across diverse *TP63* mutations.

While this study provides compelling evidence that p63β can compensate for p63α loss in the epidermis and offers a promising therapeutic strategy for AEC syndrome, there are limitations. The functional redundancy between p63α and p63β appears to be restricted to the epidermis. Developmental defects observed in the palate and limbs of p63α-deficient mice underscore the requirement of α-specific domains for morphogenesis in non-cutaneous tissues, limiting the scope of isoform switching as a universal therapeutic strategy. Although CRISPR-mediated exon 13 deletion effectively shifts isoform usage in patient-derived keratinocytes, the efficiency and safety of this approach remain untested *in vivo*. The risk of off-target editing and potential immunogenicity of Cas9-based systems must be rigorously evaluated before clinical application. In conclusion, our study reveals that p63β can functionally compensate for the loss of p63α in the epidermis, and that exon 13 deletion restores keratinocyte fitness in AEC syndrome.

## Materials and Methods

### Generation of mouse models

Conditional p63Δ13 knockout mice (p63Δ13^fl/fl^) were generated by inserting LoxP sites flanking exon 13 of the *Trp63* gene, together with a PGK-neomycin (PGK-neo) selection cassette flanked by FRT sites (Suppl. Figure 1A). To generate the P63-3xFlag conditional knock-out construct (pL253-P63Lox-3xFlag) we inserted a LoxP site 632bp upstream of exon 13 in pL253-P63Lox1-Ex14-3xFlag plasmid (Russo et al., 2018). Briefly, miniarms (PCR product: EmP63U-EmP63L and FmP63U-FmP63L) were cloned into NotI-EcoRI (EmP63U/L) or BamHI-SalI (FmP63U/L) sites in the pL451 vector (Liu, Jenkins et al., 2003). The “armed” Frt-PGK-Neo-Ftr-LoxP box and pL253-P63Lox1-Ex14-3xFlag plasmid were co-electroporated into the recombinogenic bacterial strain EL350 to obtain the final plasmid. This construct was targeted to the endogenous p63 locus in E14TG2a (129/Ola) embryonic stem (ES) cells through homologous recombination. Twenty-four hours after electroporation ES cells were selected with 100 μg/ml G418 (ThermoFisher Scientific, cat.no. 10131035) for 7 days. Neomycin-resistant ES clones were screened at 5’ and 3’ for the correct insertion in the *Trp63* endogenous locus by PCR analysis using the TaKaRa LA Taq® DNA Polymerase (Takara, cat.no. RR002B) with an oligonucleotide annealing in the neomycin cassette and in the genomic DNA (5’Arm Forward and Reverse; 3’Arm Forward and Reverse). An ES positive clone was injected into C57BL/6 blastocysts at Biogem S.c.a r.l. animal facility (Italy, https://www.biogem.it/index.php/en/). The obtained chimeric mice were tested for germline transmission by breeding with C57BL/6 females. The neomycin cassette was removed by breeding the first generation of heterozygous mice with transgenic mice carrying the Flip recombinase (Russo et al., 2018). p63Δ13^fl/fl^ mice were crossed with CAG-Cre mice (Tg(CAG-cre)1Nagy) (Nagy, 2000) to obtain CAG-Cre; p63Δ13^fl/fl^ mice. K14-Cre (Krt14-CreΔneo) knock-in mice (Huelsken et al., 2001) were used to induce p63 exon 13 depletion in stratified epithelia at late stages of embryonic development. Oligonucleotide primers are in (Suppl. Table 1). Mice were housed in individually ventilated cages under controlled temperature (22–24°C) and humidity (45– 65%) with a 12-hour light/dark cycle. Food and water were provided ad libitum. All animal protocols described here were carried out according to the European Community Council Directive 86/609/EEC, upon approval by the CEINGE Biotecnologie Avanzate Ethical Committee for Animal Experimentation and by the Italian Ministry of Health Italian Ministry of Health (D. Lgs 116/92, auth. 3/12/12, and D5A89.58 auth. 928/2021-PR).

### Cell Culture

Primary keratinocytes were isolated from the skin of P2 neonatal mice. Skin samples were incubated overnight at 4°C, floating on a Dispase II solution (0.80 U/mL; ThermoFisher Scientific cat.no. 17105041) in HBSS supplemented with 10 mM HEPES, 0.075% sodium bicarbonate, and antibiotic/antimycotic (ThermoFisher Scientific, cat.no. 15240062). The following day, epidermal sheets were separated from the dermis using forceps and incubated in 1 mL of Accutase (Euroclone, cat.no. ECB3056D) for 20 minutes at room temperature. Subsequently, 500 µL of 0.25% Trypsin–EDTA (ThermoFisher Scientific, cat.no. 25200072) was added for 4 minutes to enhance enzymatic dissociation. Trypsin activity was neutralized by the addition of 10% Fetal Bovine Serum (FBS;) (Cha1115L, Cytivia Hyclone) in HBSS. The resulting single-cell suspension was filtered through a 70 µm cell strainer (Corning, cat.no. 431751) to remove undigested tissue and debris. The isolated keratinocytes were seeded into collagen-coated dishes and cultured for 6 days at 34°C and 8% CO_2_ in a low-calcium medium (0.05 mM CaCl_2_) supplemented with 4% calcium-depleted characterized FBS and Epidermal Growth Factor (Corning, cat.no. 354052) as previously described (Russo et al., 2018).

HEK293T, HDFn, BJ-HDF and 3T3-J2 feeder cells (Rheinwald & Green, 1975) cell lines were grown in DMEM, 10% FBS, 1% L-Glutamine (Euroclone, cat.no. ECB3000), 1% Penicillin-Streptomycin (Euroclone, cat.no. ECB3001D), at 37°C and 5% CO_2_. Primary human keratinocytes were plated (3000 cells/cm^2^) on 80% confluent mitotically blocked 3T3-J2 feeder cells, and grown at 37°C, 5% CO_2_ in humidified atmosphere in DMEM (ThermoFisher Scientific, cat. no. 61965-026,) and Ham’s F12 medium (ThermoFisher Scientific, cat.no. 31765-027) (2:1 mixture) containing HyClone characterized U.S. origin FBS (CytivaLifeSCience, cat.no. SH30071.03), 1% Penicillin-Streptomycin, 4mM L-Glutamine, 0.18 mM Adenine (Sigma-Aldrich, cat.no. A2786), 5 mg/mL Transferrin (Sigma-Aldrich cat.no. T8158), 5 mg/mL Insulin (ThermoFisher Scientific cat.no. 12585014), 0.1 nM cholera toxin (Sigma-Aldrich, cat.no. C8052), 0.4 mg/mL hydrocortisone (Millipore, cat.no. 386698), 2 nM triiodothyronine (Sigma-Aldrich, cat.no. T6397), 10 ng/mL epidermal growth factor, and 10 µM ROCK inhibitor Y27632 (Enzo Life Sciences, cat.no. ALX-270-333).

### Retroviral, lentiviral and adenoviral preparation

Retroviruses were produced in HEK293T cells by transient co-transfection of pBABE-ΔNp63α, pBABE-ΔNp63β constructs or pMXs-KLF4 plasmid (pMXs-Klf4 was a gift from Shinya Yamanaka; Addgene plasmid # 13370) and amphotropic viral envelope plasmid (pAmpho) in the presence of Lipofectamine 2000 (ThermoFisher Scientific, cat. no. 11668027) according to the manufacturer’s instructions. Cell supernatants containing the retroviruses were collected 48 hours after transfection, then fresh medium was added and collected again 72 hours after transfection. The retrovirus preparation was filtered using 0.45 μm filters to remove cellular debris. Two days after infection, cells were selected with 1 μg/ml of puromycin for 3 days. Mouse primary keratinocytes were infected 4 days after plating with adenovirus carrying GFP as control or Cre recombinase (a kind gift of Okairos AG) at MOI 100 for 2 hr in low calcium medium without serum and EGF. Cells were collected two days after infection.

### Fibroblast to keratinocyte-like cell conversion

HDFn and HDF-BJ cells were co-transduced twice at 30% confluency with retroviruses encoding either p63α or p63β, in combination with KLF4, in the presence of 8 µg/mL polybrene (Sigma-Aldrich, cat.no. TR-1003-G). Two days post-transduction, cells were trypsinized and replated at 20% confluency in medium supplemented with 2 µg/mL puromycin (ThermoFisher Scientific, cat.no. A1113903) for 4 days to select for successfully transduced cells. Following antibiotic selection, cells were maintained in DMEM supplemented with 10% fetal bovine serum (FBS) for an additional 15 days in the absence of puromycin. The culture medium was replaced every two days. On day 15, cells were either fixed for immunofluorescence or harvested for downstream analyses. Quantitative assessment of conversion efficiency was performed by immunofluorescence using anti-KRT5 and anti-KRT14 antibodies. Results were validated by RT-qPCR and western blot analysis for KRT14.

### RNA extraction, RT-qPCR and RNA-seq

Total RNA was extracted from mouse or human keratinocytes using the Direct-zol MicroPrep Kit (Zymo Research, cat.no. R2061), or from mouse epidermis using the RNeasy Plus Mini Kit (Qiagen, cat.no. 74134). For RT-qPCR, cDNA was synthesized using the LunaScript RT SuperMix (New England Biolabs, cat.no. E3010) and relative mRNA expression was assessed using the Luna Universal qPCR Master Mix (New England Biolabs, cat.no. M3003) on an Applied Biosystems 7500 Real-Time PCR System. Data were analyzed using the 2^−ΔΔCT^ method (Schmittgen & Livak, 2008). mRNA expression levels were normalized to Actin for mouse samples and to RPLP0 for human cells. Oligonucleotide primers used are described in Suppl Table 1.

For RNA-seq analysis of embryonic epidermis, total RNA was extracted from the epidermis of E18.5 embryos carrying either the CAG-Cre; p63Δ13^fl/fl^ or p63Δ13^fl/fl^ genotype. RNA concentration was measured using the Qubit 2.0 Fluorometer (ThermoFisher Scientific). Libraries were prepared from 125 ng of total RNA using the 3’DGE mRNA-seq digital mRNA-sequencing service (NEGEDIA; https://negedia.com/en/) (Xiong, Soumillon et al., 2017), which included library preparation, quality assessment and sequencing on a NovaSeq 6000 sequencing system (Illumina Inc.) using a single-end, 100 cycle strategy. The raw data were analyzed by NEGEDIA digital mRNA-seq pipeline (v1.0) involving quality filtering, trimming, alignment to the reference genome and counting by gene (Anders, Pyl et al., 2015, Bushnell, 2014, Dobin, Davis et al., 2013). Differential expression analysis was performed using edgeR5 (Robinson, McCarthy et al., 2010). Genes having an average count of < 10 cpm and an FDR≥ 0.05 were filtered out. For RNA-seq analysis of primary keratinocytes, total RNA was extracted from the mouse keratinocytes isolated from P2 mice carrying either the K14-Cre; p63Δ13^fl/fl^ or p63Δ13^fl/fl^ genotype and cultured in supplemented Low Calcium medium for 7 days. Samples were processed as described above.

For mRNA-seq in human keratinocytes, Nextflex Rapid Directional RNA-Seq kit 2.0 with Poly(A) 2.0 (PerkinElmer) was performed using NovaSeq sequencing system (Illumina Inc.) using a 2x150 bp 30M paired-end run (Procomcure NGS, Austria; https://ngs.procomcure.com). Fastq files containing reads were trimmed using Trim Galore! to remove sequences with low quality and adapter. Alignment was performed with RNA STAR on mouse mm10 reference assembly. The expression levels of genes were determined with featureCounts (Liao, Smyth et al., 2014), and the differential expression analysis was performed using DESeq2 (Dobin et al., 2013). Genes were filtered using the FDR < 0.05, log_2_FC≥0.5 or log_2_FC≤-0.5, and average counts ≥10. Splicing analysis was performed importing the bam files into the IGV software for coverage and splicing visualization (Robinson, Thorvaldsdottir et al., 2023).

### ChIP-seq

Mouse dorsal skin was harvested P0, and the epidermis was separated from the dermis by overnight incubation in Dispase solution at 4 °C and single-cell suspensions were obtained by incubation in Accutase for 20 minutes at room temperature. Cells were cross-linked in 1% formaldehyde for 10 minutes at 34 °C. After quenching with 125 mM glycine for 5 minutes at room temperature, cells were collected by centrifugation and washed twice with ice-cold PBS containing 1 mM PMSF and a protease inhibitor cocktail (Roche, cat.no. 11697498001). Pellets containing 4 × 10⁶ cells were resuspended in 500 µL of SDS lysis buffer (50 mM Tris-HCl pH 8.1, 1% SDS, 10 mM EDTA) supplemented with protease inhibitors, incubated on ice for 10 minutes. ChIP-seq was performed as previously described (Antonini et al., 2015) with the following modifications. Samples were sonicated in 250 µL aliquots using a Bioruptor® Plus (Diagenode) at high power (60 s ON, 90 s OFF) for 45 minutes at 4 °C to obtain chromatin fragments of approximately 300 bp. Immunoprecipitation was performed on 4 mg of total protein using 4 µg of anti-p63 antibody (clone EPR5701, Abcam, cat.no. 124762) and 2.1 mg of Dynabeads Protein A/G (1:1 ratio; ThermoFisher, cat.no. 10006 and 10007).

Purified DNA was used for library construction using the KAPA Hyper prep kit (Roche; according to manufacturer’s protocol). The prepared libraries were then sequenced on the NextSeq 500, according to Illumina protocols. Two independent samples were processed for each condition, and Fastq files of wild type and p63α knock-out were mapped to the Mus musculus genome (mm10 build) using Bowtie2 (Langmead & Salzberg, 2012). Call peaks from alignment results were performed using MACS2 (2.1.1.20160309) (Feng, Liu et al., 2012) on pooled replicates, combining the two independent samples per condition. Highly stringent conditions were selected (FDR40, Fold enrichment 10, pile-up100). Heat map, profile plot and clustering were obtained using the chromatin analysis and exploration tool ChAsE (Younesy, Nielsen et al., 2016), using the summit ± 2.5Kb of the peaks (-LOG10FDR ≥ 40, Fold enrichment ≥ 10, pile-up ≥100) obtained from p63 mutant ChIP-seq as reference bed file, and the p63 wild type and mutant bigwig files. Clustering was performed using K-means (k=3) GREAT, version 4.0.4) (McLean, Bristor et al., 2010), with the following parameters: “basal plus extension” mode, assigning peaks to genes with a proximal window of 5 kb upstream and 1 kb downstream of the transcription start site, and a distal extension up to 100 kb.

Peaks associated with both p63α and p63β (cluster1 and cluster2 Figure 3) or predominantly associated with p63β (cluster 3) were annotated to nearby genes based on the mm10 assembly from the UCSC Genome Browser. Genomic locations were assigned using the Genomic Regions Enrichment of Annotations Tool (GREAT, version 4.0.4) (McLean et al., 2010), with the following parameters: “basal plus extension” mode, assigning peaks to genes with a proximal window of 5 kb upstream and 1 kb downstream of the transcription start site, and a distal extension up to 100 kb. Enriched GO biological processes of p63 target genes were identify using Metascape (Zhou, Zhou et al., 2019). To identify the consensus for transcription factors associated with the identify genes Enrichr (https://maayanlab.cloud/Enrichr/) was used with a combination of ENCODE datasets and ChEA3 (Keenan, Torre et al., 2019). Motif discovery and enrichment analysis was performed using XSTREME (v.5.5.7) using the top 1000 genomic regions bound by p63 according to the FDR using default parameters. p63 binding motifs and their locations in the sequences was identified using CentriMo with 500 bp sequences centered on the summit of each peak (Bailey, Johnson et al., 2015). To evaluate the regulatory potential of differentially bound genomic regions, we assessed their overlap with an Expanded Registry of mouse cCRE registry from the ENCODE project (Moore, Pratt et al., 2024). Bedtools intersect (Galaxy; https://usegalaxy.org) was used to compute the fraction of regions that overlapped ENCODE cCREs by at least one base pair. The analysis was performed separately for all differentially enriched regions, and for individual clusters identified by k-means clustering (e.g., cluster 3, Figure 3D). Overlap percentages were calculated as the number of overlapping regions divided by the total number of input regions.

### Blue native (BN)-PAGE, SDS-PAGE, and Western blot

For Blue Native-PAGE (BN-PAGE), cells were scraped on ice in native lysis buffer (25 mM Tris-HCl pH 7.5, 150 mM NaCl, 10 mM MgCl₂, 10 mM EDTA pH 8.0, 20 mM CHAPS, 2 mM DTT) supplemented with protease and phosphatase inhibitors, and incubated for 1 hour on ice in the presence of benzonase (Merck Millipore, cat. no. 70746). Clarified lysates were mixed with 20% glycerol and 5 mM Coomassie G-250 (ThermoFisher Scientific, cat. no. BN2004), and loaded onto 3–12% Novex Bis-Tris native gradient gels (ThermoFisher Scientific, cat. no. BN1003BOX), following the manufacturer’s protocol. After electrophoresis, proteins were transferred to PVDF membranes and processed for western blotting using standard procedures. For SDS-PAGE, cells were lysed directly in 2× SDS sample buffer (10% glycerol, 0.01% bromophenol blue, 0.0625 M Tris-HCl pH 6.8, 3% SDS, 5% β-mercaptoethanol) supplemented with protease and phosphatase inhibitors, boiled, and resolved on standard denaturing SDS-PAGE gels. Proteins were transferred to PVDF membranes for western blotting. Membranes were blocked in PBS containing 0.2% Tween-20 and 5% non-fat dry milk, and incubated with primary antibodies either for 2 hours at room temperature or overnight at 4 °C. After washing, membranes were incubated with horseradish peroxidase (HRP)-conjugated secondary antibodies for 1 hour at room temperature. Signal detection was performed using Clarity Max Western ECL substrate (Bio-Rad, cat. no. 1705062) and visualized with the ChemiDoc Gel Imaging System (Bio-Rad). Densitometric analysis was conducted using Image Lab Software v6.1.0 (Bio-Rad). The list of primary and secondary antibodies used in this study is provided in Suppl. Table 2.

### Histology and Immunostaining

Dorsal skin, embryonic heads, or whole embryos were dissected, fixed in 4% paraformaldehyde (Electron Microscopy Sciences, cat.no.157-4) in PBS, and embedded in paraffin. Sections of 5– 7 µm thickness were prepared for histological or immunofluorescence analysis. Hematoxylin and eosin (H&E) staining or H&E and Alcian blue staining were performed following standard procedures. For immunostaining of paraffin sections, antigen retrieval was performed by heating samples in 0.01 M citrate buffer (pH 6.0) using a microwave oven. For immunofluorescence staining, sections were blocked in 1% BSA/0.02% Tween/PBS for 1 h at room temperature.

Primary antibodies were incubated o/n at 4 °C in blocking buffer and washed in 0.2% Tween/PBS. Secondary antibodies were incubated at room temperature for 1 h and washed in 0.2% Tween/PBS, and 4,6-diamidino-2-phenylindole (DAPI) was used to identify nuclei. Fluorescence signals were visualized using a Zeiss Axioskop2 Plus microscope and a Zeiss Apotome 2.0. Macroscopic images were acquired using a Leica M205FA stereomicroscope. Zeiss Axioscan Z1 was used for Hematoxylin Eosin staining. For immunostaining of cultured cells, cells were fixed in 4% PFA for 10 minutes at room temperature, washed three times with PBS, and permeabilized with 0.5% Triton X-100 (Merck, cat.no. T8787) for 5 minutes. After three additional PBS washes, cells were blocked with 10% goat serum (ThermoFisher Scientific, cat.no. 16210072) and incubated overnight at 4 °C with primary antibodies diluted in PBS containing 2% goat serum. Following PBS washes, cells were incubated with appropriate secondary antibodies and Hoechst 33342 nuclear stain (ThermoFisher Scientific, cat.no. H1399) for 45 minutes at room temperature. A complete list of primary and secondary antibodies used in this study is provided in Suppl. Table 2.

### Alizarin and Alcian staining

For skeletal preparations, embryos were eviscerated and the limbs dissected and skinned. Dissected limbs were fixed in 90% ethanol for 7 days at room temperature, followed by fixation in 100% acetone for an additional 2 days. Cartilage staining was performed by incubating specimens in Alcian Blue solution (2 mg/mL Alcian Blue in 80% ethanol and 20% glacial acetic acid) for 3 days. Following staining, specimens were rehydrated through sequential incubation in 70%, 40%, and 15% ethanol, and then rinsed in distilled water. Next, specimens were incubated in 1% potassium hydroxide (KOH) for 4 hours, and then subjected to bone staining in Alizarin Red S solution (1 mg/mL Alizarin Red S in 1% KOH) for 2 days. After staining, samples were further incubated in 1% KOH for multiple washes and then cleared by sequential incubation in increasing concentrations of glycerol in KOH: 0.8% KOH/20% glycerol, 0.5% KOH/50% glycerol, and 0.2% KOH/80% glycerol, for 3–5 days each. Fully cleared specimens were photographed and stored in 100% glycerol for long-term preservation.

### Skin Barrier Assay and TEWL Measurement

Skin permeability was assessed using a toluidine blue dye penetration assay. Embryos were dehydrated by sequential immersion in 25%, 50%, 75%, and 100% methanol (2 minutes each), followed by rehydration through the reverse gradient (75%, 50%, 25% methanol) and a final wash in PBS prior to staining. Then embryos were incubated in filtered toluidine blue aqueous solution (0.0125% for E17.5 and 0.1% for E18.5) for 10 minutes at 4 °C. Following staining, embryos were rinsed thoroughly in PBS and imaged using a Leica M205FA stereomicroscope. TEWL was measured using a Tewameter (TM NANO; Courage and Khazaka) on the dorsal skin at P0.

### Excisional skin-wound-healing assay

Two circular, full-thickness excisional wounds (6 mm in diameter) were created on the lower dorsal skin of each animal, approximately 2 cm apart, one on each side of the midline. Procedures were performed under general anesthesia. At the time of sample collection (day 3 and day 7 post-wounding), animals were euthanized and skin samples encompassing the wound area were excised post-mortem. Tissues were fixed overnight at 4 °C in buffered 4% formaldehyde solution, then embedded in paraffin. Paraffin blocks were sectioned at 7 µm thickness for histological analysis. Morphometric evaluation was performed on H&E-stained sections taken from the center of day 3 (D3) and day 7 (D7) wounds. Quantitative analysis included: (i) wound closure, (ii) distance between wound edge hair follicles (HF) as a measure of tissue contraction, and (iii) length of the newly formed epithelium to assess re-epithelialization. All measurements were carried out using ImageJ software. The wound area was also monitored macroscopically by imaging wounds daily from day 0 to day 10 post-surgery. Wound size at each time point was normalized to the original wound area measured on day 0 for each individual wound.

### EdU Incorporation assay

To assess proliferation, primary human keratinocytes were pulsed with 1 mM of EdU (ThermoFisher Scientific, cat.no. C10640) in keratinocyte growth medium for 3 h at 37°C before fixation. Cells were rinsed twice with 1X PBS and fixed with warm 4% paraformaldehyde for 10 minutes at room temperature. EdU was revealed by Click-iT reaction accordingly to manufacturer instructions.

### Genome editing in human keratinocytes

CRISPR/Cas9 mediated genome editing was performed as previously described (Bamundo, Palumbo et al., 2024). Briefly, synthetic guide RNAs (gRNAs) targeting sequences flanking exon 13 of the *TP63* gene were designed using the Custom Alt-R™ CRISPR-Cas9 guide RNA design tool (Integrated DNA Technologies, IDT) (Suppl. Table 1). Ribonucleoprotein (RNP) complexes were assembled using 10^4^ pmol of Alt-R™ S.p. HiFi Cas9 Nuclease V3 (IDT, cat.no. 1081058) per reaction. Two independent RNP reaction mixes were prepared: RNP1 contained 120 pmol of sgRNA targeting the region upstream of exon 13 and Alt-R™ S.p. HiFi Cas9 protein; RNP2 contained 120 pmol of sgRNA targeting the region downstream of exon 13 and Cas9 protein. Each RNP complex was assembled separately by incubating sgRNA with Cas9 protein for 20 minutes at room temperature. Following individual complex formation, RNP1 and RNP2 were combined and incubated together for an additional 5 minutes at room temperature before transfection. A total of 2 × 10⁵ human keratinocytes were resuspended in 20 µL of supplemented primary cell nucleofection solution (P3 Primary Cell 4D-Nucleofector™ X Kit; Lonza, cat.no. V4XP-3032) and mixed with the pre-formed RNP complexes along with 4 mM Cas9 electroporation enhancer (Alt-R™ Cas9 Electroporation Enhancer; IDT, cat.no. 1075916). Cells were electroporated using the 4D-Nucleofector™ System (Lonza) with the DS-138 program. Nucleofection efficiency was assessed by PCR and western blot analysis.

### Dispase-based keratinocyte dissociation assay

Keratinocytes were seeded at a density of 6,000 cells/cm² onto mitotically inactivated 3T3-J2 feeder cells in six-well plates and cultured until they reached full confluence, typically after ∼15 days. At confluence, the resulting epidermal sheets were gently washed with DPBS containing Ca²⁺ and Mg²⁺, then incubated with 2.4 U/mL Dispase II (Roche, cat.no. 0494207801) for 30 minutes at 37 °C. Following detachment, the intact monolayer was carefully transferred into a 15 mL Falcon tube containing 5 mL of PBS with Ca²⁺ and Mg²⁺ and subjected to mechanical stress by manually inverting the tube 50–100 times. Fragmentation of the sheet was used as a measure of cell-cell adhesion strength as previously described (Ferone et al., 2013).

### Colony-Forming Efficiency (CFE) Assay

The Colony-Forming Efficiency (CFE) assay was performed to assess the clonogenic potential of basal keratinocytes. A total of 1,000 NHEK-p63WT and NHEK-p63Δ13 cells were seeded on mitotically inactivated 3T3-J2 feeder layers and cultured under standard conditions as described above. After 12 days of culture, the entire well surface was imaged using the CellDiscoverer7 automated imaging system (Zeiss) with oblique illumination. Colonies were counted, and CFE values were calculated using the following formula: CFE (%) = (Number of Colonies / Number of Cells Plated) × 100. Colony size was also quantified using Zen Blue software (Zeiss, version 3.2).

### Population doubling capacity

The proliferative potential of keratinocytes was evaluated by assessing their population doubling capacity (PDC) over serial passages. NHEK-p63WT and NHEK-p63Δ13 cells were continuously cultured on mitotically inactivated 3T3-J2 feeder layers for up to ten passages. At each passage, cells were trypsinized, counted using trypan blue exclusion to assess viability, and reseeded at a density of 3,000 cells/cm² for the subsequent passage. Population doublings were calculated at each passage using the formula: PD = 3.322 × log(N / N₀), where N represents the number of viable cells harvested at the end of the passage, and N₀ is the number of clonogenic cells initially seeded.

### Statistics

Data analysis, statistical testing, and visualization were performed using Prism (GraphPad Software, San Diego, CA, USA; www.graphpad.com). The specific statistical tests applied are detailed in the corresponding figure legends. All experiments were independently repeated at least three times using biologically distinct samples. For details regarding the statistical analysis of transcriptomic data, please refer to the corresponding paragraph in the Methods section.

## Data availability

RNA-seq and ChIP seq raw data generated in this study have been deposited in GEO (GSE297765, GSE297766, GSE297767, GSE298875). For detection of the splicing isoforms in mouse embryogenesis the following data were used: GSE97213 for surface ectoderm and GSE150702 for limb ectoderm.

## Acknowledgments.

We are grateful to Dr. Diego Di Bernardo (TIGEM, Naples) for his extensive support and expertise in bioinformatics analysis, and to Dr. Davide Cacchiarelli (TIGEM, Naples) for his insightful guidance on transcriptomic analyses and for providing BJ-HDF cells. We thank Lucille Lopez-Delisle (University of Geneva) for her help in generating Sashimi plots of mouse limb data. We acknowledge the CEINGE–Franco Salvatore Advanced Light Microscopy Facility for support with cell imaging and analysis. We also thank Jo Zhou and Joost H. A. Martens at the RIMLS-Science Sequencing Facility, Radboud University, for ChIP-seq library preparation and sequencing, and Jo Zhou for her continued scientific feedback. This work was supported by the Telethon Foundation (grant GGP20124) and the Italian Association for Cancer Research (AIRC, grant IG 25116). This work is dedicated to the memory of Dr. Daniel Aberdam—a generous colleague and dear friend.

## Notes

### Competing Interest Statement

The authors have declared no competing interest.

