## Supplementary data for "Targeted p63 isoform switch corrects dominant mutations in AEC syndrome without disrupting epidermal homeostasis"

### Supplementary Figure and Figure Legends

Suppl. Figure 1

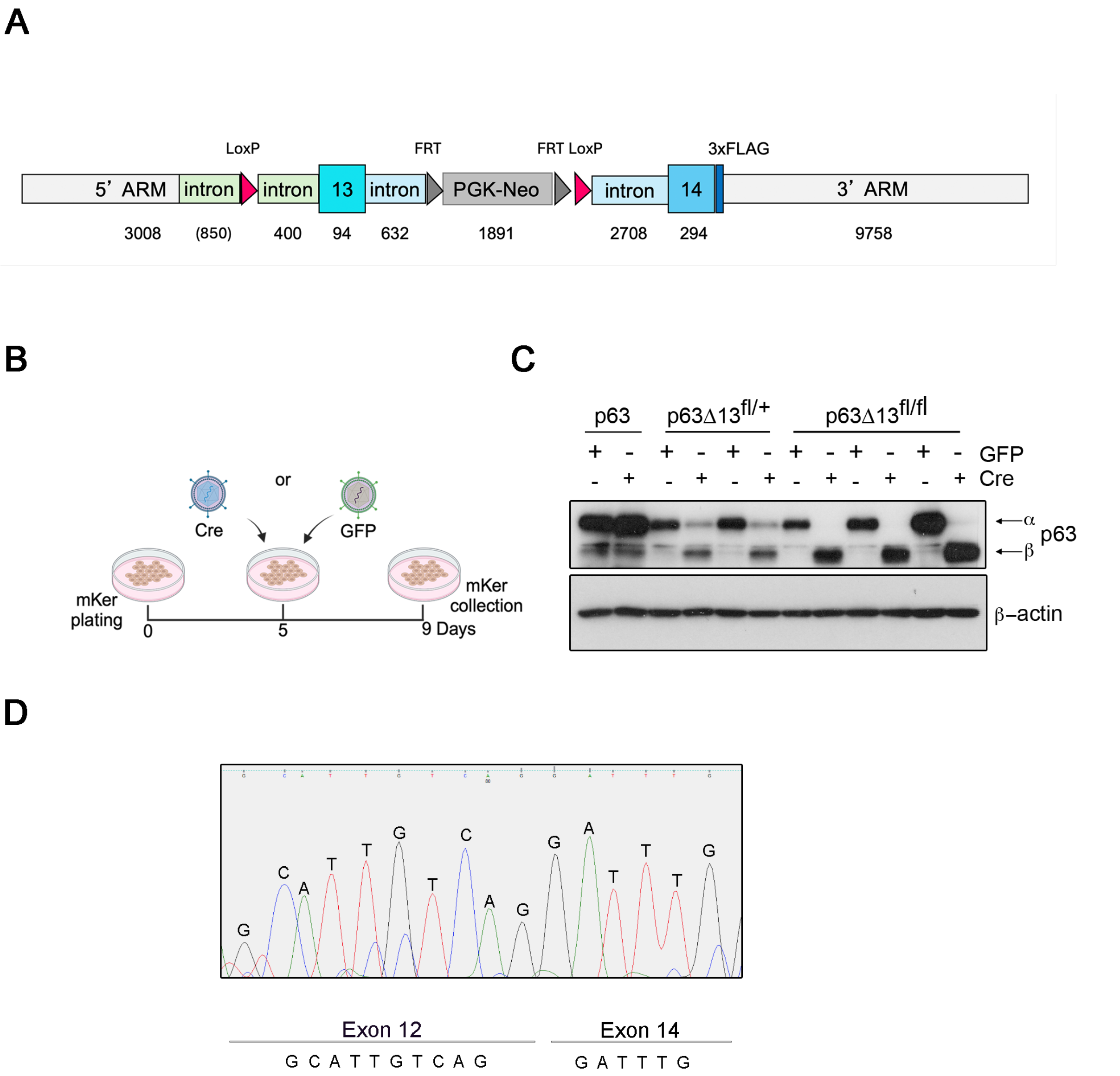


**Suppl. Figure 1. Generation and molecular characterization of p63α-deficient mice and primary keratinocytes.**

A. Scheme of the targeting construct used to generate p63α-specific isoform knock-out. The construct includes a 5′ homology arm (5′ ARM; 3008 bp) containing part of intron 12, followed by a LoxP site upstream of exon 13 (94 bp). Exon 13 is flanked by intronic regions (400 bp and 632 bp), and a PGK-Neo selection cassette (1891 bp) flanked by FRT sites. A second LoxP site was inserted downstream, followed by the remaining 3′ portion of intron 13 (2708 bp), the coding region of exon 14 (294 bp) tagged with 3×FLAG, and the 3′ homology arm (3′

ARM; 9758 bp), which includes part of the 3′ untranslated region (2798 bp; not indicated). The PGK-Neo cassette was removed by Flp recombinase. Segment lengths are indicated in bp.

B. Experimental scheme showing transduction of mouse primary keratinocytes (mKer) with adenoviruses expressing either Cre recombinase or GFP control. Created with BioRender.com.

C. Western blot showing p63α and p63β protein expression levels in transduced mKer. Actin was used to normalize samples.

D. Electropherogram confirming proper junction between exon 12 and exon 14 in cDNA derived from CAG-Cre; p63Δ13$^{fl/fl}$ epidermis.

Suppl. Figure 2

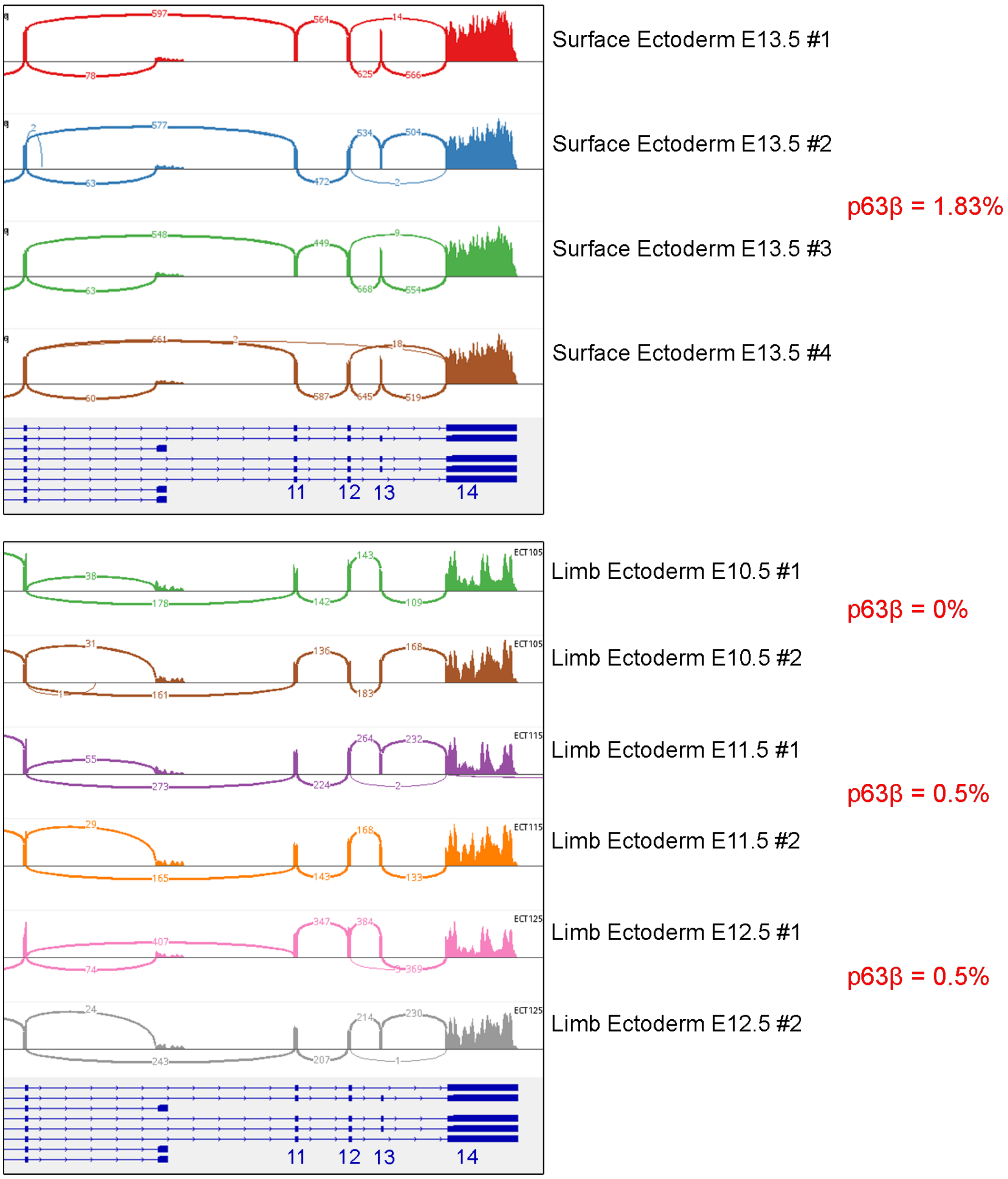


**Suppl. Figure 2. Alternative splicing of *Trp63* in surface and limb ectoderm during embryogenesis.**

Sashimi plots of RNA-seq data from surface ectoderm (E13; *n*=4) and limb ectoderm (E10.5–E12.5; *n*=2 per stage) reveal very low expression levels of the p63β isoform. Arcs represent splice junction reads; arc thickness corresponds to the number of reads connecting the two exons indicated.

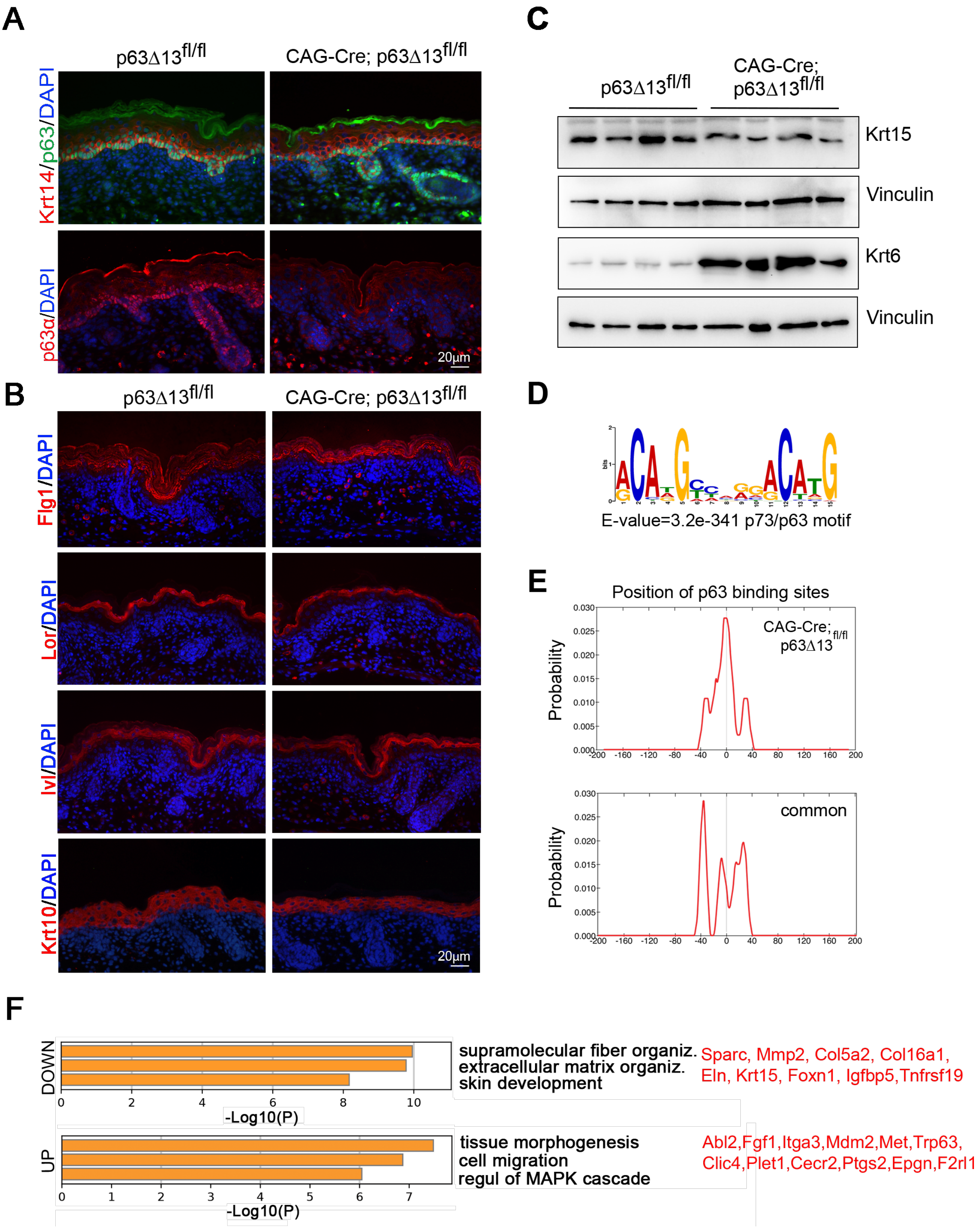


**Suppl. Figure 3. p63β compensates for p63α loss with minimal effects on epidermal differentiation and chromatin binding profiles.**

A. Representative immunofluorescence analysis of P0 epidermis using a p63α-specific (red) and a pan-p63 (green) antibodies. p63α expression is absent in CAG-Cre; p63Δ13$^{fl/fl}$ epidermis, while pan-p63 staining confirms expression in the basal layer, consistent with p63β compensation. Krt14 (red) marks basal keratinocytes. Nuclei are stained with DAPI (blue). Scale bar: 20 µm.

B. Representative immunostaining for late differentiation markers shows comparable expression of filaggrin (Flg), loricrin (Lor), involucrin (Ivl), and keratin 10 (Krt10) in control and mutant P0 epidermis. Nuclei are stained with DAPI (blue). Scale bar: 20 µm.

C. Western blot analysis of Krt15 and Krt6 in P0 epidermal extracts. Krt15 expression is reduced, while Krt6 is ectopically induced in mutant samples. Vinculin was used to normalize samples (n=4 mice/genotype).

D. *De novo* motif discovery in the top 1,000 p63β-specific binding regions reveals enrichment for the canonical p63/p73 motif.

E. Distribution of canonical p63 binding motifs (DRCATGTCNNRACAYGYM) in mutant-specific (top; cluster 3) and shared (bottom; clusters 1–2) regions. Shared sites display broader motif distribution, suggesting higher redundancy.

F. Gene Ontology (GO) enrichment analysis of genes upregulated (bottom) or downregulated (top) in CAG-Cre; p63Δ13$^{fl/fl}$ epidermis versus controls.

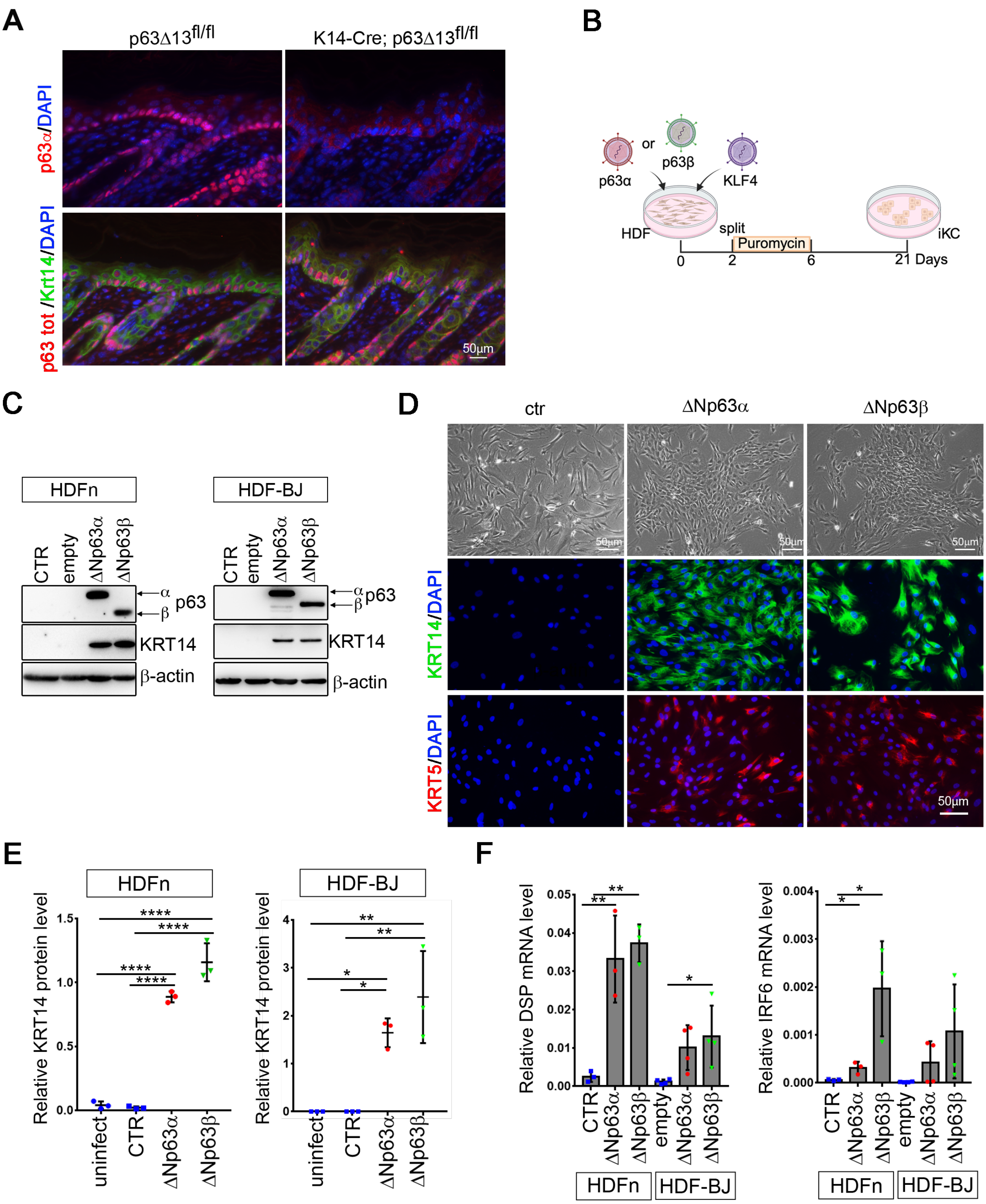


**Suppl. Figure 4. Induction of epidermal commitment in dermal fibroblasts by overexpressing p63β.**

A. Representative immunofluorescence staining of p63Δ13$^{fl/fl}$ and CAG-Cre; p63Δ13$^{fl/fl}$ skin at P5. Sections were stained with a p63α-specific antibody (red, upper panels) or a pan-p63 antibody (red) together with Keratin 14 (Krt14, green, bottom panels). Nuclei were counterstained with DAPI (blue). Scale bar: 50 µm.

B. Experimental scheme: HDF were transduced with retroviruses carrying p63α and KLF4 or p63β and KLF4, and analysed 18 days post-transduction. Both primary and the HDF-BJ cell line were used with similar results. Created with BioRender.com.

C. Western blot analysis of KRT14 expression in transduced HDFs (HDFn and BJ). Actin served as a loading control. Actin was used to normalize samples.

D. Representative bright field images (top), and immunofluorescence for KRT14 (middle) and KRT15 (bottom) in transduced HDFs. Nuclei were counterstained with DAPI (blue). Scale bar: 50 µm.

E. Densitometric quantification of KRT14 protein levels normalized to Vinculin, corresponding to panel C ($n = 3$ independent experiments).

F. RT-qPCR analysis of DSP and IRF6 mRNA expression following p63α + KLF4 or p63β + KLF4 transduction ($n \geq 3$ independent experiments).

Suppl. Figure 5

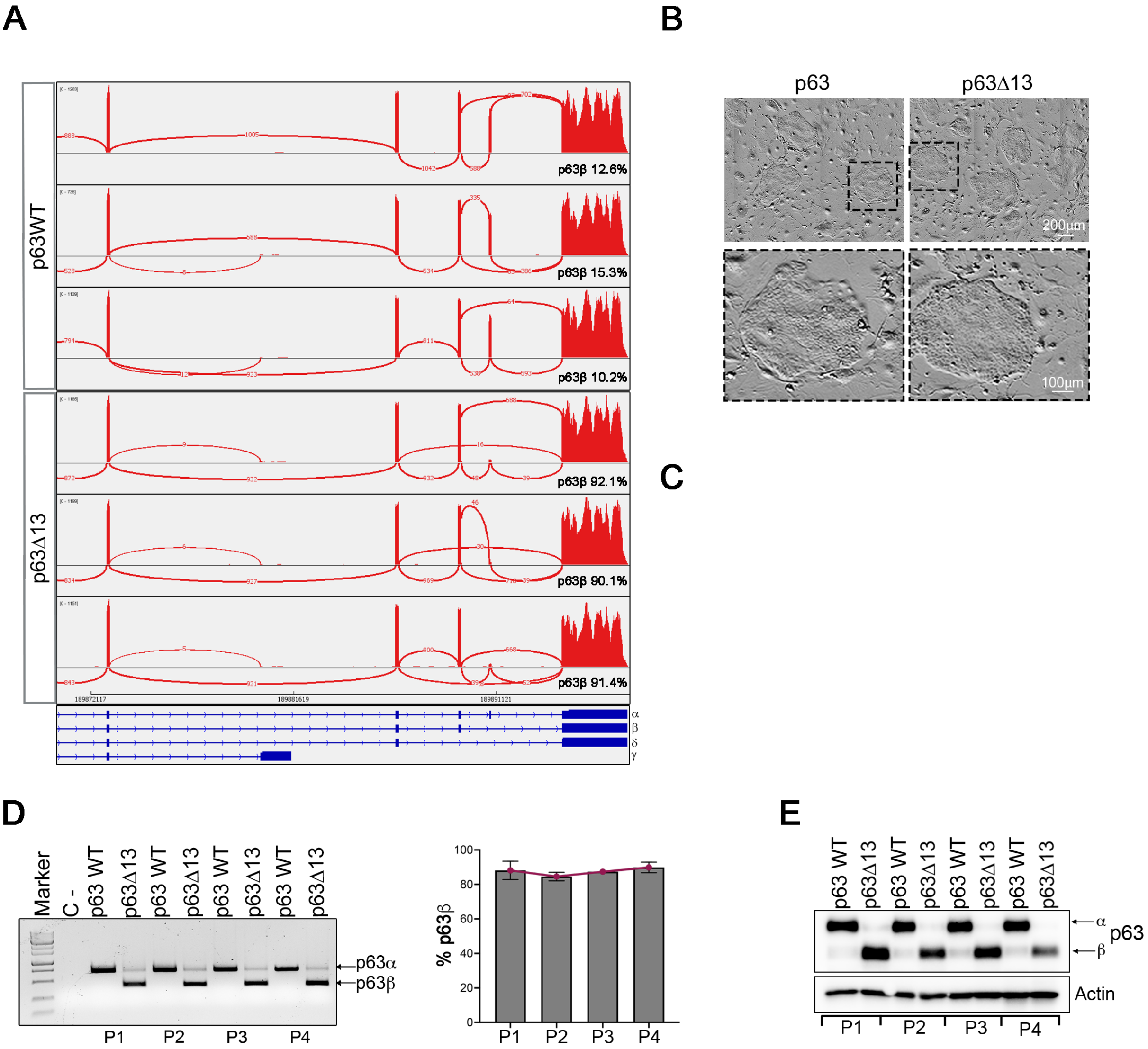


**Suppl. Figure 5. p63β expressing primary human newborn keratinocytes retain their properties despite several passages *in vitro*.**

A. Sashimi plots of RNA-seq data showing splicing between exons 11 and 14 in control (p63WT) and CRISPR/Cas9-edited (p63Δ13) keratinocytes. Arcs represent splice junction reads; arc thickness correlates with read count.

B. Representative bright-field images of p63WT and p63Δ13 keratinocytes at low (top) and high (bottom) magnification. Scale bars: 200 µm (top), 100 µm (bottom).

C. Experimental scheme: keratinocytes were nucleofected with RNPs, then split every 6 days at a density of 3,000 cells/cm². Cells were collected at each passage (P1–P4) for analysis. Created with BioRender.com.

D. PCR analysis of genomic DNA from p63WT and p63Δ13 keratinocytes at different passages (P1–P4) showing stable exon 13 deletion. Right: densitometric quantification of p63β expression (% of total) across four independent experiments.

E. Western blot analysis of ΔNp63α and ΔNp63β protein levels at passages P1–P4 in p63WT and p63Δ13 keratinocytes. Actin was used to normalize samples. (n=4 independent experiments).

Suppl. Figure 6

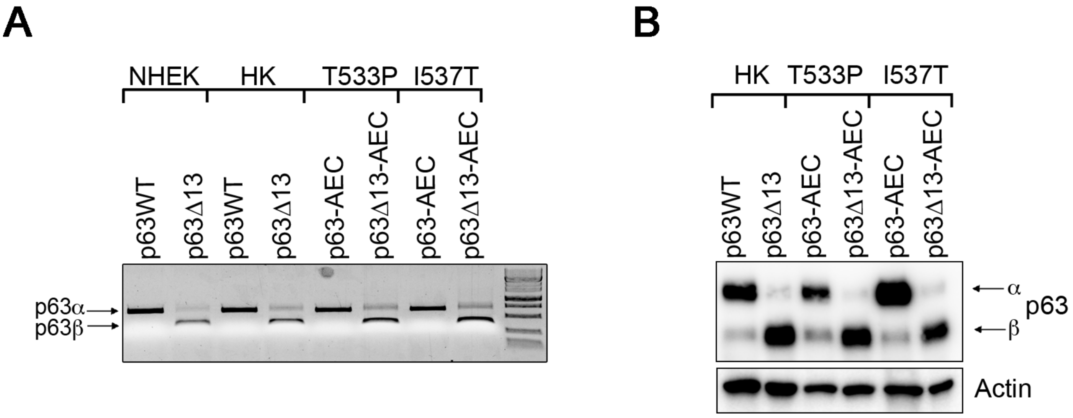


**Suppl. Figure 6. Validation of exon 13 deletion in wild-type and AEC-mutant keratinocytes.**

A. PCR analysis of genomic DNA from wild-type (p63WT), AEC-mutant (T533P and I537T), and corresponding exon 13-deleted keratinocytes (p63Δ13 and p63Δ13-AEC) showing exon 13 deletion.

B. Western blot analysis showing ΔNp63α and ΔNp63β protein expression in wild-type and AEC-mutant keratinocytes with or without exon 13 deletion. Actin was used to normalize samples. (n=4 independent experiments).
